# Nuclear GSK-3β and Oncogenic KRas Promote Expansion of Terminal Duct Cells and the Development of Intraductal Papillary Mucinous Neoplasm

**DOI:** 10.1101/2020.07.14.202952

**Authors:** Li Ding, Kaely Roeck, Cheng Zhang, Brooke Zidek, Esther Rodman, Yasmin Genevieve Hernandez-Barco, Jin-San Zhang, William R. Bamlet, Ann L. Oberg, Lizhi Zhang, Nabeel Bardeesy, Hu Li, Daniel D. Billadeau

**Affiliations:** Division of Oncology Research, College of Medicine, Mayo Clinic, Rochester, MN 55905, USA; Department of Molecular and Experimental Therapeutics, College of Medicine, Mayo Clinic, Rochester, MN 55905, USA; Center for Cancer Research, Harvard Medical School, Boston, MA 02115, USA; Center for Precision Medicine, The First Affiliated Hospital of Wenzhou Medical University; Institute of Life Science, Wenzhou University, Zhejiang, China; Department of Health Sciences Research, College of Medicine, Mayo Clinic, Rochester, MN 55905, USA; Department of Laboratory Medicine and Pathology, College of Medicine, Mayo Clinic, Rochester, MN 55905, USA

**Keywords:** GSK-3β, Oncogenic KRas, Pancreatic Cancer, Intraductal Papillary Mucinous Neoplasm (IPMN), terminal ducts

## Abstract

Intraductal papillary mucinous neoplasm (IPMN) represents one type of pancreatic ductal adenocarcinoma (PDA) precursor lesion, however its cell-of-origin remains unclear. Here we describe a new mouse model in which pancreas-specific Cre activation of a nuclear glycogen synthase kinase-3β transgene is combined with oncogenic KRas (referred to as KNGC). KNGC mice show accumulation of neoplastic ductal cells at 4-weeks that progressively develop into IPMN with low-grade dysplasia in advanced age. RNA-sequencing identified expression of several terminal duct cell lineage genes including Agr2 and Aqp5. Interestingly, Aqp5, a water channel, was found to be required for the development of IPMN lesions in KNGC mice. Staining of human IPMN samples indicates that these preneoplastic lesions also arise from expansion of the terminal duct population. Altogether, these data highlight the utility of the KNGC model for understanding the biology of IPMN and potential utility in defining predictive biomarkers of IPMN – PDA development.

**Statement of significance:** Understanding the cell-of-origin of IPMN is crucial to developing early detection methods that specifically target aggressive precursors of PDA. This work, using a novel mouse model, identifies Aqp5-modulated development of Agr2^+^ terminal ducts that could potentially serve as a clinical biomarker for IPMN.

## Introduction

Pancreatic ductal adenocarcinoma (PDA) is predicted to be the second leading cause of cancer-related deaths in the USA by 2030. The 5-year relative survival rate of all stages combined is less than 10%, with 80% of patients having a locally advanced tumor at diagnosis, which highlights the significant need for the development of novel early-detection strategies and models to understand PDA tumor development (1–4).

Three major types of morphologically distinct PDA precursors have been identified to date, including pancreatic intraepithelial neoplasia (PanIN), intraductal papillary mucinous neoplasm (IPMN) and mucinous cystic neoplasm (5), each likely have an unique impact on disease biology, response to therapy and prognosis (6,7). Although there has been an increase in understanding of PDA disease progression, established from well appreciated studies of patient samples and the development of genetically engineered mouse models (GEMM) (8–10), the cell-of-origin of PDA remains controversial.

Sequencing from human specimens has shown an overlap of somatic mutations between PanIN and IPMN, including the activating mutation of KRas, the most frequent and earliest genetic alteration in PDA (11,12), and the secondary loss-of-function mutation of tumor suppressors TP53 and SMAD4/DPC4, particularly in high-grade lesions (13). Moreover, lineage-tracing experiments and cell-type specific activation of oncogenic KRas coupled to other genetic alterations have been conducted and the collective data demonstrate that, although acinar cells possess the highest plasticity to be transformed and are responsible for most PanIN and IPMN lesions through acinar-to-ductal metaplasia (ADM), ductal cells harboring oncogenic KRas can also give rise to precursor lesions sharing similar histology features, and might also serve as a cell-of-origin for PDA (9,10,14–16). Thus, delineating the origin of PDA precursor lesions with deeper molecular insight could facilitate the development of clinical tools for early-detection and improve PDA patient outcomes in the clinic.

Glycogen synthase kinases, GSK-3α and GSK-3β, were originally identified as key enzymes regulating glycogen metabolism (17–19). However, accumulating evidence has suggested a role for these kinases in several human malignancies (20–22) and has identified them as therapeutic targets in pancreatic cancer (23). Significantly, GSK-3β is overexpressed in PDA and aberrant nuclear accumulation, as well as increased mRNA expression of GSK-3β, has been associated with high-grade tumors (24,25). Moreover, our prior studies showed that mutant KRas increases GSK-3β gene expression *in vivo* and *in vitro* (26,27). However, the studies conducted so far to address the multifaceted role of GSK-3β in pancreatic cancer *in vivo* have mostly relied on loss-of-function approaches including knockout mice or GSK-3 inhibitors (23). The role of nuclear GSK-3β in pancreatic cancer progression *in vivo*, particularly as it pertains to PDA precursor lesion development, is unknown.

To address this, we generated a novel pancreas-specific nuclear GSK-3β expression and oncogenic KRas activation model (KNGC) to investigate the function of nuclear GSK-3β and its impact on PDA precursor lesion initiation and progression. Unexpectedly, we observed profound pancreatic cyst development and expansion of ductal cells in 4-week old KNGC mice and subsequent low-grade IPMN lesions in KNGC mice of advanced age. RNA-sequencing data indicated that nuclear GSK-3β and oncogenic KRas reprogrammed pancreatic progenitor cells toward the ductal lineage, resulting in a loss of acinar cells and the expansion of ductal cells expressing anterior gradient 2 (Agr2) and Aquaporin 5 (Aqp5). Agr2 and Aqp5 expression was found to be associated with the terminal or intercalated ductal cell compartment. Aqp5 knockout in KNGC mice dramatically abrogated differentiation and development of Agr2^+^ terminal ducts. Using tissue microarrays, we show that human IPMN lesions also arise from Agr2^+^/Aqp5^+^ terminal ducts. Taken together, these results highlight pancreatic terminal duct cells as the likely cell-of-origin for human IPMN, and provide a novel mouse model for biomarker discovery, IPMN development and progression to PDA.

## Results

### GSK-3β ablation limits oncogenic KRas induced pancreatic cancer development

A genetically engineered mouse model, expressing activated mutant KRas (G12D) in the pancreas, develops pancreatic cancer with long latency (28), which can be substantially accelerated by experimentally inducing chronic pancreatitis or crossbreeding with mutations/deletions of the tumor suppressors TP53 or Ink4a/Arf (29,30). We have previously demonstrated that GSK-3β expression is a target of activated KRas signaling in pancreatic cancer cells and pancreas specific deletion of GSK-3β in KRas mutant mice reduces cearulein-induced ADM and PanIN lesion development, while abrogating KRas mutation-dependent proliferation (26,27). To examine the impact of GSK-3β ablation on pancreatic cancer progression, the pancreas was collected from 8-to 10-month old Pdx1-cre/LSL-KRasG12D (KC) and Pdx1-cre/LSL-KRasG12D/GSK-3β^F/F^ (RKO) littermates (Figure 1A). The total number of ductal lesions and their grade were scored in representative pancreatic sections. Depletion of GSK-3β was confirmed by immunofluorescent staining of GSK-3β and the pancreatic ductal cell marker CK19. Strong GSK-3β signal was detected in CK19^+^ neoplastic ducts from KC mice but not RKO mice (Figure 1B). Consistent with a previous report (28), the majority of pancreas tissue from 9-month old KC mice were replaced by metaplastic ducts and surrounded by desmoplasia (Figure 1C). In contrast, RKO mice had diminished numbers of ADM and low-grade PanIN lesions (1 and 2), as well as a reduced incidence of high-grade PanIN-3 and invasive cancer (Figure 1C and D) compared to KC mice. Since GSK-3β has been shown to participate in the regulation of pancreatic cancer cell proliferation (23,27), we examined the proliferation of neoplastic ducts in KC and RKO mice. As shown in Figure 1E and F, we observed an accumulation of EdU^+^/CK19^+^ ductal cells within neoplastic areas in KC mice, which were dramatically reduced in RKO mice. Taken together, these data suggest that GSK-3β is required for oncogenic KRas-driven cell proliferation and deletion of GSK-3β impairs preneoplastic lesion and pancreatic cancer development in KC mice.

**Figure 1.**
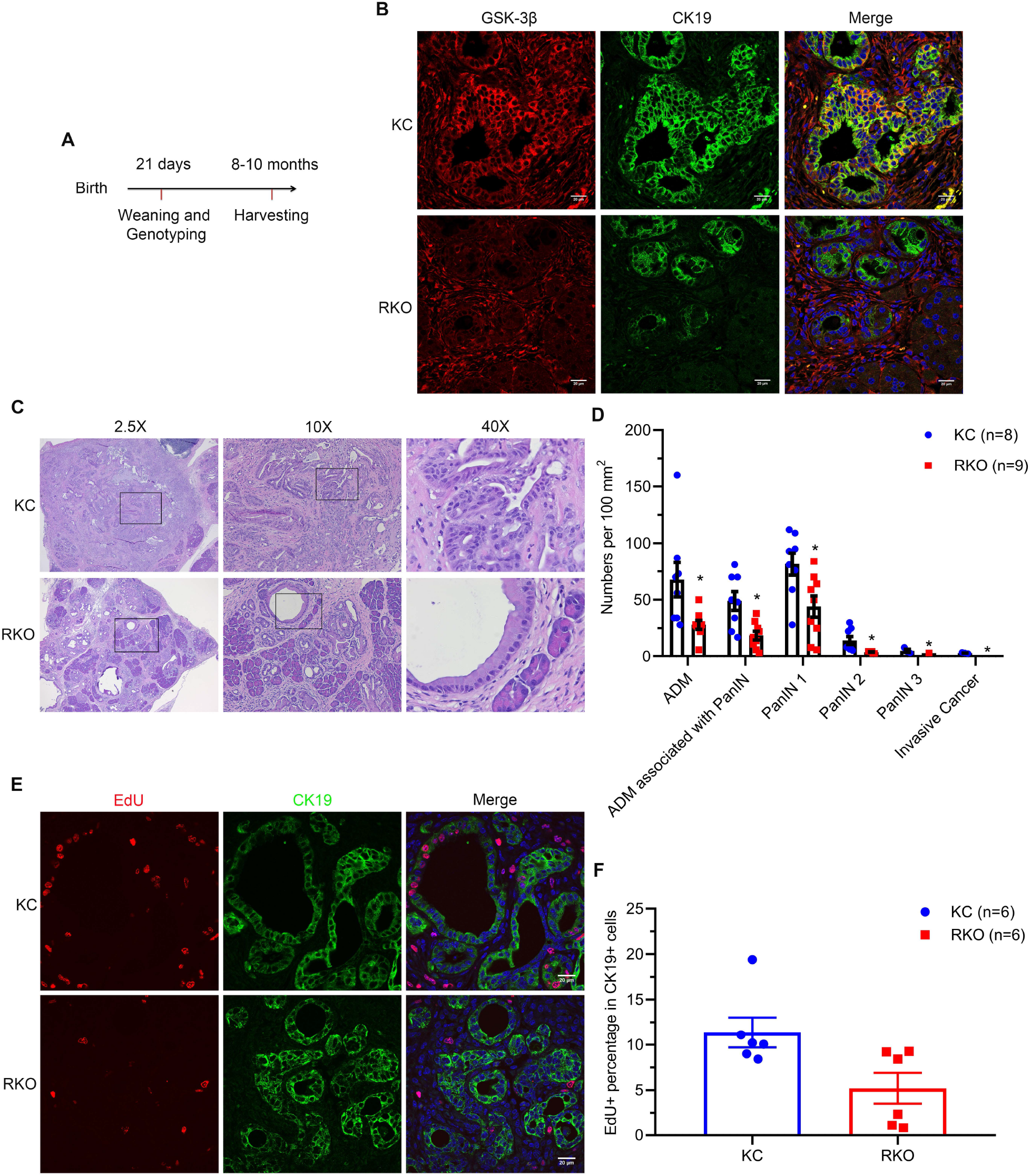
GSK-3β deletion limits oncogenic KRas-induced pancreatic cancer development. (A) Scheme for aging-induced pancreatic cancer progression model and analysis. (B) Immunofluorescence staining of GSK-3β (red) and CK19 (green) from pancreatic sections of aging KC and RKO mice. Nuclei were counter-stained with Hoechst (blue). (C) H&E stained pancreatic sections from KC and RKO mice. Black boxes indicate magnified area. (D) H&E stained tissue samples from KC and RKO mice were evaluated and quantitatively analyzed for numbers per 100 mm^2^. Data were analyzed and expressed as mean ± SEM. *P<0.05 RKO versus KC mice. (E) Double-labeling of pancreatic sections from aging KC and RKO mice was performed using incorporated EdU detection kit (red) and CK19 (green) antibodies. Nuclei were counterstained with Hoechst (blue). (F) Quantification of the percentage of EdU positive neoplastic ductal cells in aging KC and RKO mice. Data were analyzed and expressed as mean ± SEM. n = 6. *P<0.05 RKO versus KC mice.

### Nuclear GSK-3β and oncogenic KRas promote the pancreatic ductal cells expansion and IPMN development

We have previously shown that GSK-3β is overexpressed in PDA and becomes localized to the nucleus in high-grade tumors (24). To characterize the role of nuclear GSK-3β in pancreatic cancer development, we generated a novel pancreas-specific nuclear GSK-3β expression model by inserting a GSK-3β transgene containing an HA tag and a nuclear localization signal sequence into the *Rosa26* locus. The transgene also contained a 5’ Lox-STOP-Lox (LSL) cassette for tissues-specific activation. Mice carrying LSL-nuclear GSK-3β (NG) were crossbred with KC mice to produce KNGC mice (Figure 2A). As shown in Figure 2B, protein extracts from both NGC and KNGC animals showed a unique HA band which was not seen in WT or KC mice. Phosphorylation of Erk1/2, a downstream target of activated KRas, was only slightly upregulated in KC mice due to limited formation of ADM and PanIN precursor lesions at this age (28). However, KNGC mice had a dramatic increase of phospho-Erk1/2, suggesting a hyper-activation of KRas signaling induced by expression of nuclear GSK-3β. Consistent with the immunoblot results, gross pathological examination revealed hallmarks of pancreatic neoplastic transformation, including visible cysts and desmoplasia in KNGC mice, but no gross pathological changes were found in the pancreas from WT, NGC and KC mice (Figure 2C).

**Figure 2.**
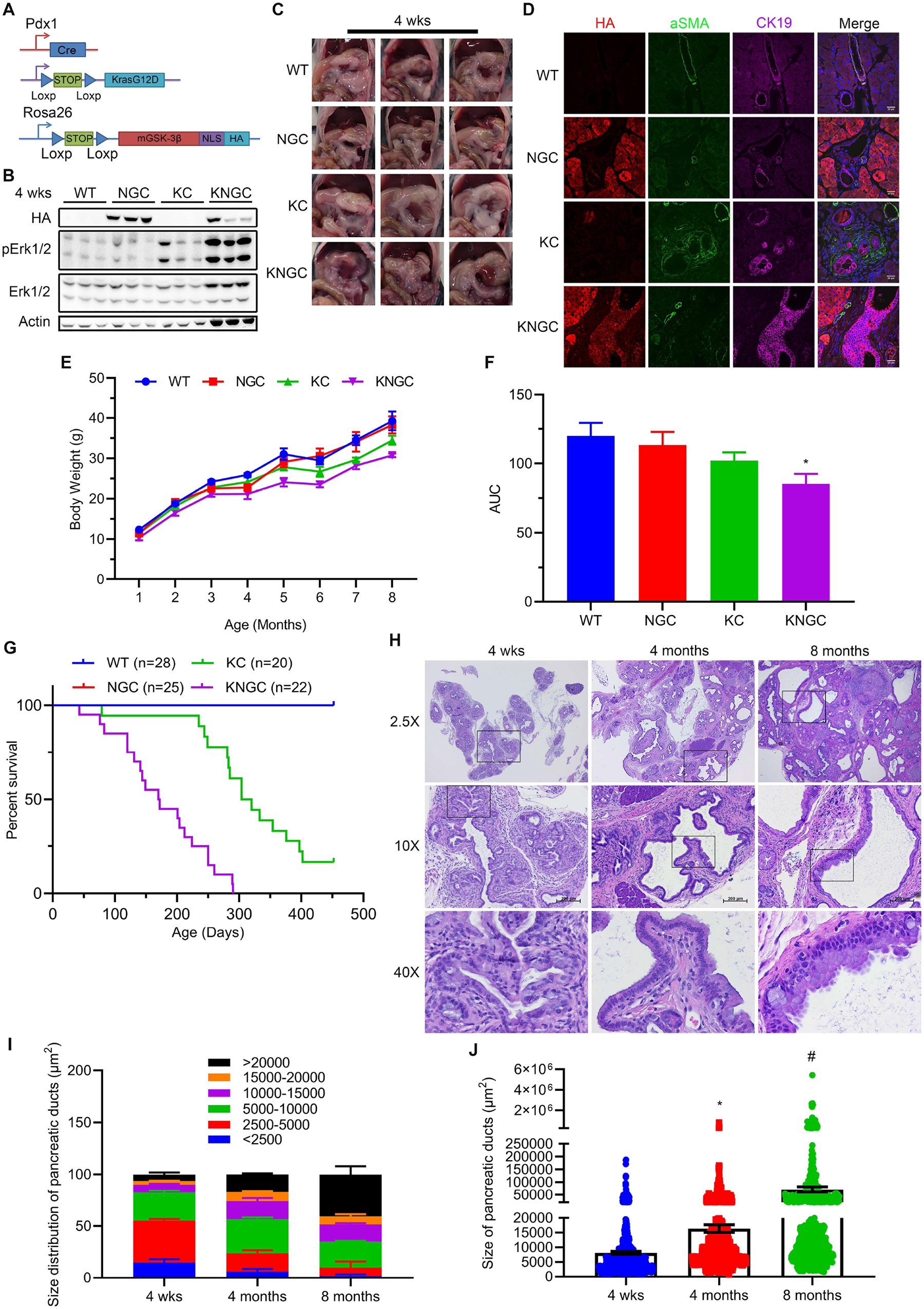
Nuclear GSK-3β and oncogenic KRas promote pancreatic ductal cell expansion and IPMN development. (A) Schematic representation of KNGC (LSL-KRasG12D/Rosa26-LSL-nuclear GSK-3β/Pdx1-Cre) mouse model. Cre expression is driven by the Pdx1 promoter-Cre transgene. Blue triangles indicate loxP sites. (B) Cell lysates from the indicated genotypes at 4 weeks of age were prepared and probed with the indicated antibodies. β-actin was used as a loading control. Shown are representative results from 6 experiments. (C) Gross pathology of pancreas and adjacent tissues from transgenic mice of indicated genotypes at 4 weeks of age. (D) Immunofluorescence staining of HA-tag (red), aSMA (green) and CK19 (purple) from pancreatic sections of the indicated genotypes at 4 weeks of age. Nuclei were counter-stained with Hoechst (blue). Body weight (E) and area under the curve (AUC) (F) of WT, NGC, KC and KNGC mice fed normal chow diet were measured at the indicated time points. Data were analyzed and expressed as mean ± SEM. At least 10 mice were included at each point. *P<0.05 KNGC mice versus the other genotypes. (G) Kaplan-Meier survival curve of the indicated genotypes. (H) H&E stained pancreatic sections from KNGC mice of 4 weeks, 4 months and 8 months. Black boxes indicate magnified area. Quantification of pancreatic duct size distribution (I) and distribution of pancreatic duct size (J) for KNGC mice were analyzed and expressed as mean ± SEM. n = 5. *P<0.05 4 months versus 4 weeks KNGC mice. #P<0.05 8 months versus 4 months KNGC mice.

Immunofluorescent staining confirmed that the HA-tag was widely expressed in acinar and hyperplastic ductal cells within the pancreas of NGC and KNGC mice, which was absent from WT and KC mice (Supplemental Fig. S1). Alpha-smooth muscle actin (αSMA), a marker for activated myofibroblasts, was associated with areas of metaplasia in KC mice but not with the areas of ductal hyperplasia observed in the KNGC mice (Figure 2D). Due to apparent loss of the normal physiologic function of the pancreas, KNGC mice had a significantly lower body weight across their lifespan and a dramatically shortened median survival of approximately 6 months as compared with KC mice (Figure 2E, F and G). Histologically, normal acinar cell clusters were rarely seen in KNGC mice. At 4 weeks of age, KNGC mice developed multifocal ductal structures and atypical ductal cells across the pancreas, which rapidly expanded with age (Figure 2H, I and J). Loss of ductal cell polarity, papillary architecture with increased nuclear/cytoplasmic ratio, and mucinous epithelial neoplasm were most frequently seen in 8-month old KNGC mice, representing the histological features of low-grade IPMN. Consistent with the pathologic definition of low-grade IPMN, no invasive cancer or metastases were observed in 8-month old age KNGC mice. Taken together, these data suggest that overexpression of nuclear GSK-3β with oncogenic KRas results in the development of low-grade IPMN lesions, a well-known precursor to PDA.

### Nuclear GSK-3β and oncogenic KRas initiate ductal hyperplasia at an early stage of pancreatic development

It is of interest that NGC mice were indistinguishable from WT littermates in every age group (data not shown), suggesting that nuclear GSK-3β expression, on its own, is insufficient to drive ductal cell expansion. The induction of acute pancreatitis in KC mice leads to acinar-to-ductal metaplasia (ADM) and the generation of PanIN lesions (31). Whether nuclear GSK-3β might promote ADM or PanIN lesion formation following pancreas injury, independent of oncogenic KRas, was unclear. To test this, we utilized cearulein to induce acute pancreatitis in WT and NGC mice and collected the pancreas 7 days after cearulein treatment (Supplemental Fig S2A). Histopathological examination showed no significant differences in pancreas histology with minimal traces of pancreatitis (Supplemental Fig S2B). These results indicate that nuclear GSK-3β expression is not sufficient to promote pancreatitis-induced ductal transformation without oncogenic KRas *in vivo*.

Given the ductal hyperplasia and atypia observed in immature KNGC mice, we generated a conditional model where nuclear GSK-3β and oncogenic KRas could be induced in the whole body of adult mice using Rosa26-Cre^ERT^ (KNG-R26CreER, Figure 3A). Four-weeks after tamoxifen induction, expression of HA-tag and mutant KRas were confirmed in KNG-R26CreER mice by immunoblotting (Figure 3B). Similar to a prior report (32), activation of KRas and nuclear GSK-3β varied in mature acinar and ductal cells of KNG-R26CreER mice as determined by immunofluorescent staining for HA (Figure 3C). Additionally, very few ADM lesions were observed in either K-R26CreER or KNG-R26CreER mice (Figure 3C). Since over 95% of the mature pancreas is comprised of acinar cells, and the expression of HA was relatively low in mature ductal cells (Figure 3C), we next sought to examine whether nuclear GSK-3β expression could increase the susceptibility of pancreatic ductal cells to oncogenic KRas-initiated tumorigenesis using Krt19-Cre^ERT^ (KNG-Krt19CreER, Figure 3D). Surprisingly, we did not observe significant ductal metaplasia in the pancreas in either K-Krt19CreER or KNG-Krt19CreER mice 2 months following tamoxifen induction (Figure 3E). These data suggest that metaplasia induced by nuclear GSK-3β and oncogenic KRas does not occur in acinar or ductal cells in the adult pancreas.

**Figure 3.**
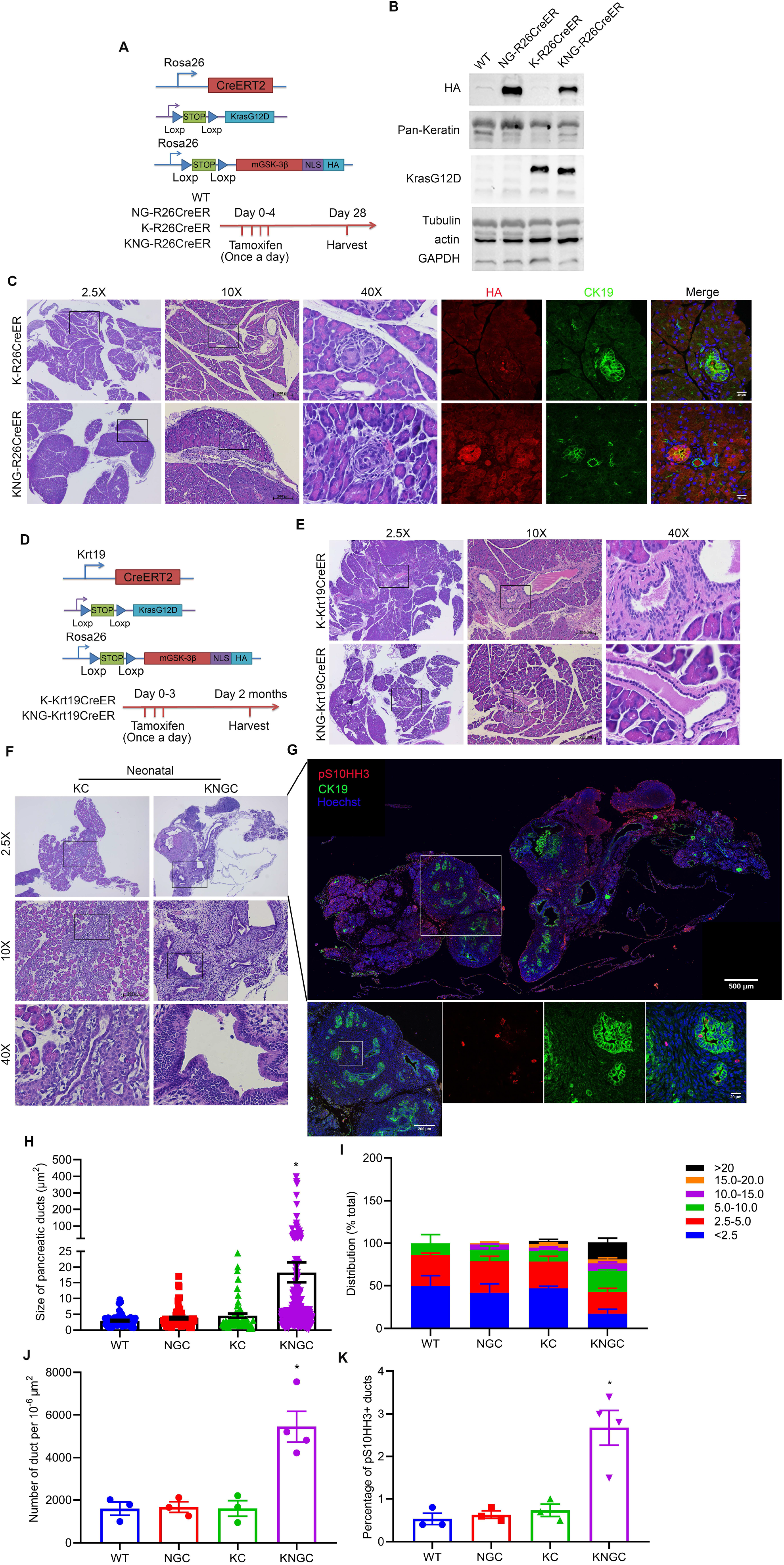
Nuclear GSK-3β and oncogenic KRas initiate ductal hyperplasia at an early stage of pancreas development. (A) Schematic representation of NG-, K- or KNG-R26CreER (LSL-KRasG12D/Rosa26-LSL-nuclearGSK-3β/Rosa26-Cre^ERT^) mouse model. Cre expression under the Rosa26 promoter was induced by tamoxifen injection in 6-8-week old mice. Blue triangles indicate loxP sites. (B) Cell lysates from pancreas of indicated genotypes 28 days post tamoxifen injection were prepared and probed with the indicated antibodies. Tubulin, β-actin and GAPDH were used as a loading control. Shown are representative results from 5 experiments. (C) H&E and immunofluorescence staining of HA-tag (red) and CK19 (green) pancreatic sections from K-R26CreER and KNG-R26CreER mice treated as above. Black boxes indicated area magnified. Nuclei were counter-stained with Hoechst (blue). (D) Schematic representation of K- or KNG-Krt19CreER (LSL-KRasG12D/Rosa26-LSL-nuclear GSK-3β/Krt19-Cre^ERT^) mouse model. Cre expression under the Krt19 promoter was induced by tamoxifen injection in 6-8-week old mice. Blue triangles indicate loxP sites. (E) H&E stained pancreatic sections from K-Krt19CreER and KNG-Krt19CreER mice treated as above. Black boxes indicate magnified area. (F) H&E stained pancreatic sections from neonatal KC and KNGC mice. Black boxes indicate magnified area. (G) Pancreas section of neonatal KNGC mice in (F) were stained for pS10HH3 (red) and CK19 (green). White boxes indicate magnified area. Nuclei were counter-stained with Hoechst (blue). Quantification of average pancreatic duct size (H), pancreatic duct size distribution (I) as well as number of ducts per 10^-4^ μm^2^ (J) and percentage of pS10HH3 positive ducts (K) were analyzed and expressed as mean ± SEM. n = 4. *P<0.05 KNGC mice versus the other genotypes.

Expression of *PDX-1/IPF1* is believed to be the earliest identifiable marker in pancreatic progenitor cells (28,33,34). Therefore, we examined the pancreata of neonatal KNGC mice, where we observed an increased number of ducts, ductal hyperplasia and enlargement, and marked increase in the number of phospho-ser10 histone H3 positive (pS10HH3+) mitotic cells (Figure 3F-K and Supplemental Fig S2C). Taken together, these data suggest that the combined expression of nuclear GSK-3β and oncogenic KRas promotes ductal cell expansion and proliferation very early in development, but is unable to do so in adult tissues.

### Transcriptional regulation of pancreatic ductal neoplasm by overexpression of nuclear GSK-3β and oncogenic KrasG12D activation

To gain insight into the cell of origin and transcriptional pathways promoting the ductal hyperplasia seen in KNGC mice, we isolated RNA from the pancreas of 4-week old WT, NGC, KC and KNGC mice and performed RNA-Seq. Consistent with the unusual composition of the pancreas in KNGC mice, we detected over 9000 significantly changed genes compared to the other genotypes (Supplemental Figure S3A). KEGG signaling pathway analysis found over 1000 pathways to be significantly activated, while 100 pathways were substantially inactivated in KNGC mice as compared to KC or NGC mice (using log2FC>2 or log2FC<-2 as a cutoff, respectively; Supplemental Figure S3B). Among the signaling pathways increased in KNGC mice were those involved in differentiation, proliferation and development (Figure 4A). In contrast, genes with decreased expression were closely associated with secretion and digestion, granule formation and cellular metabolism (Figure 4B). As can been seen in Figure 4C, markers for acinar cells, normal ductal cells, human PDA precursors, and multiple subsets of pancreatic cancer associated fibroblasts (Pan-CAF) (35–38), were significantly changed in KNGC mice compared to the other genotypes (complete list can be found in Supplemental Excel Table S3). The differential expressions of mucin genes are associated with subtypes of IPMN (13). KNGC mice expressed a subset of mucins primarily associated with the gastric type of IPMN, including Muc5a/c and Muc6 (Figure 4D). We next validated the RNA-Seq data via quantitative PCR for two genes from each compartment (Figure 4E). In addition, significantly increased protein levels of pankeratin and Agr2, together with diminished expression of amylase, were confirmed by immunoblot (Figure 4F). Moreover, immunofluorescent staining of pancreata from KNGC mice with CK19 and amylase showed that the acinar tissue had been largely replaced with CK19^+^ hyperplastic ducts (Figure 4G and H and Supplemental Figure S3C and D).

**Figure 4.**
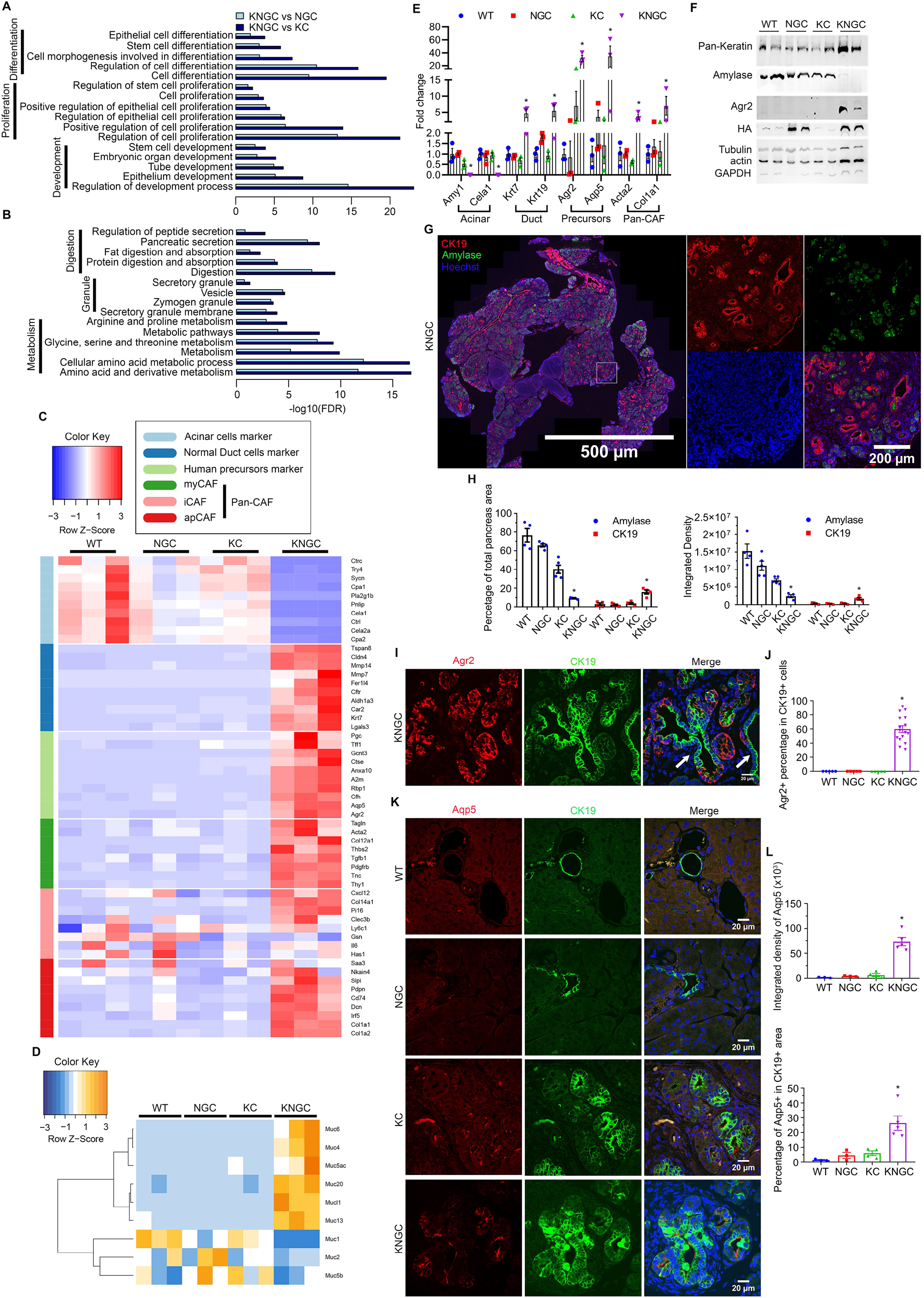
Transcriptional regulation of pancreatic ductal neoplasia by nuclear GSK-3β and oncogenic KrasG12D. (A) KEGG cellular processes enriched for genes with increased expression in KNGC mice compared with NGC or KC. (B) KEGG cellular processes enriched for genes with decreased expression in KNGC mice compared with NGC or KC. (C) and (D) Heatmap was generated for selected gene sets related to the indicated groups using normalized gene expression. Colors are assigned based on raw z-scores. (E) Real-time PCR quantification of the indicated genes from 4-week old WT, NGC, KC or KNGC mice. TBP, β-actin, RPLP0 and GAPDH were used as internal housekeeping gene controls. Data were analyzed and expressed as mean ± SEM. n = 3. *P<0.05 KNGC mice versus the other genotypes. (F) Cell lysates from pancreas of 4 week WT, NGC, KC and KNGC mice were prepared and probed with the indicated antibodies. Tubulin, β-actin and GAPDH were used as a loading control. Shown are representative results from 3 experiments. (G) Immunofluorescence staining of CK19 (red) and Amylase (green) from pancreatic sections of 4-week old KNGC mice. White boxes indicate magnified area. Nuclei were counter-stained with Hoechst (blue). (H) Quantification of percentage of total pancreas area and integrated density were analyzed and expressed as mean ± SEM. *P<0.05 KNGC mice versus the other genotypes. (I) Immunofluorescence staining of Agr2 (red) and CK19 (green) from pancreatic sections of 4-week old KNGC mice. White arrow indicated Agr2 negative/CK19 positive cells. Nuclei were counter-stained with Hoechst (blue). (J) Quantification of Agr2 positive percentage in CK19 cells was analyzed and expressed as mean ± SEM. *P<0.05 KNGC mice versus the other genotypes. (K) Immunofluorescence staining of Aqp5 (red) and CK19 (green) from pancreatic sections of 4-week old WT, NGC, KC and KNGC mice. Nuclei were counter-stained with Hoechst (blue). (L) Quantification of Aqp5 positive percentage in CK19 cells and integrated density were analyzed and expressed as mean ± SEM. *P<0.05 KNGC mice versus the other genotypes.

Two genes that were highly upregulated in KNGC mice were Agr2 and the water channel Aqp5, both of which are expressed in PDA (39,40). Therefore, we investigated the expression of Agr2 and Aqp5 in the pancreas of KNGC mice. Consistent with a previous report (41), KNGC mice showed strong Agr2 expression in close to 60% of CK19^+^ ducts, while similarly aged KC mice only expressed Agr2 in a limited number of metaplastic ducts (Figure 4I and J). Similar to Agr2 staining, Aqp5 was detected in a few early ADM lesions of KC mice, but had significantly higher distribution in CK19^+^ ducts in KNGC mice (Figure 4K and L). To determine if expression of Agr2 and Aqp5 were regulated in a GSK-3β dependent manner, we stained Agr2 and Aqp5 in aging KC and RKO mice. Interestingly, conditional GSK-3β ablation in KC mice substantially limited the development of Agr2^+^/Aqp5^+^ ducts (Supplemental Figure S3E and F). Taken together, these data suggest that expression of GSK-3β and oncogenic KRas result in a transcriptional switch from pancreatic acinar cells to neoplastic ductal cells, with expression of pancreatic tumor subtype markers.

### Expression of nuclear GSK-3β with oncogenic KRas leads to the development of two distinct ductal populations

To further characterize the neoplastic ducts arising in the KNGC model, we utilized a protocol that has been previously used to capture lectin^+^ ductal cells from the pancreas (37). This protocol combines antibody capture of ductal cells that are recognized by the lectin-binding protein *Dolichos Bifluros Agglutin* (DBA) with magnetic beads, resulting in separation of a DBA^+^/CK19^+^ ductal cell pool from a DBA^-^ cell population containing other cell types from within the pancreas (Figure 5A). Fluorescence microscopy confirmed the capture of DBA-FITC labeled cells from 4-week old KNGC mice (Supplemental Figure S4A). We observed equivalent expression of pan-Keratin in DBA^-^ and DBA^+^ ducts, with significantly higher levels of Agr2 protein in the DBA^-^ ductal pool as compared to the DBA^+^ ductal pool (Figure 5B and C). Consistent with our immunoblotting and histology data, qRT-PCR showed that DBA^-^ ducts from KNGC mice exhibited diminished acinar cell marker expression, but elevated ductal marker expression similar to the DBA^+^ ductal structures. More importantly, compared to DBA^+^ ducts, DBA^-^ ducts from 4-week old KNGC mice had increased expression of Agr2, Aqp5 and cystic fibrosis transmembrane conductance regulator (CFTR) (Figure 5D), which is another highly induced gene in KNGC mice (Figure 4C), and has been shown to colocalize with Aqp5 in human pancreatic intercalated/terminal duct cells (42). Consistent with a prior report (43), isolated DBA^-^ ductal cells from aging KC mice were enriched in acinar markers, while DBA^+^ ducts possessed much higher expression of ductal markers (Figure 5D). These results indicate that the ductal cell population expanding in KNGC mice is not only different than that found in aged KC mice, but represents a unique ductal population lacking expression of the lectin bound by DBA.

**Figure 5.**
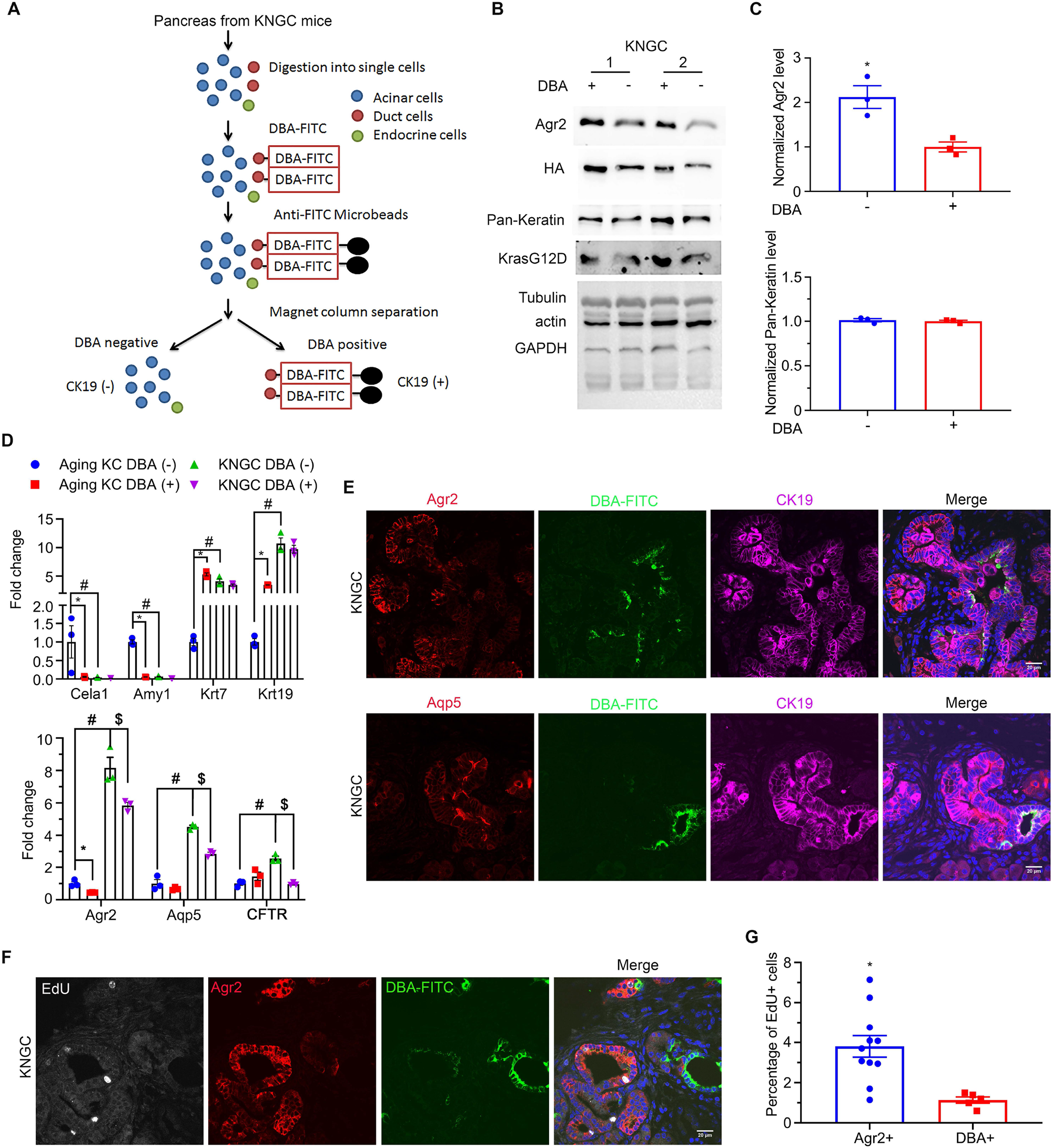
KNGC mice develop two distinct ductal populations. (A) Illustration of the experimental approach to purify lectin-expressing ductal cells that can be enriched through binding of DBA. (B) Cell lysates from isolated DBA negative and positive cells of two individual 4 week old KNGC mice were prepared and probed with the indicated antibodies. Tubulin, β-actin and GAPDH were used as a loading control. Shown are representative results from 3 experiments. (C) The average signal intensity of Agr2 and pan-Keratin was quantified and expressed as mean ± SEM. n=3. *P<0.05 DBA negative versus DBA positive samples. (D) Real-time PCR quantification of the indicated genes from isolated DBA negative and positive samples from aging KC and 4 week old KNGC mice. TBP, β-actin, RPLP0 and GAPDH were used as internal housekeeping gene controls. Data were analyzed and expressed as mean ± SEM. n = 3. *P<0.05 DBA positive versus DBA negative samples of aging KC mice. #P<0.05 DBA negative from KNGC versus DBA negative from aging KC mice. $P<0.05 DBA negative versus DBA positive samples of KNGC mice. (E) Immunofluorescence staining of Agr2 (red, upper panel) or Aqp5 (red, lower panel) with DBA-FITC (green) from pancreatic sections of 4week old KNGC mice. Nuclei were counter-stained with Hoechst (blue). (F) Immunofluorescence staining of EdU (white), Agr2 (red) and DBA-FITC (green) from pancreatic sections of 4-week old KNGC mice. Nuclei were counter-stained with Hoechst (blue). (G) Quantification of EdU positive percentage in Agr2 positive or DBA-FITC positive cells was analyzed and expressed as mean ± SEM. *P<0.05 EdU positive in Agr2 positive versus DBA-FITC positive cells.

We next examined the differential expression of Agr2 and Aqp5, along with DBA, in neoplastic ducts from KNGC mice using immunofluorescence. Co-staining of Agr2, Aqp5 and DBA-FITC showed strongly positive Agr2 and Aqp5 stained ducts that were distinct from those detected using DBA-FITC (Figure 5E). Similar staining patterns were also observed in aging KC mice (Supplemental Figure S4B). Strikingly, and consistent with the role of Agr2 in PanIN lesion initiation and PDA progression (44–46), EdU labeling showed increased DNA synthesis in Agr2^+^ ductal cells compared to DBA-FITC^+^ ductal cells (Figure 5F and G). Similar results were seen in aging KC mice, where EdU^+^ cells also accumulated in Agr2^+^ ducts. Ablation of GSK-3β in KC mice led to reduced EdU^+^ cell numbers, which was partially due to the limited proportion of ducts with Agr2 expression (Supplemental Figure S4C). Taken together, these data suggest that GSK-3β and oncogenic KRas lead to the development of a DBA^-^/Agr2^+^ ductal cell pool with more proliferative potential.

### Aqp5 is necessary for the differentiation and growth of terminal ducts in KNGC mice

The water channel Aqp5 was first described to localize mainly in terminal/intercalated and interlobular ducts in the human pancreas (42,47), also characterized as DBA^-^ (48). Analysis of our RNA-seq data indicated that Aqp5 had the highest log2FC among the 19 significantly increased genes compared between both NGC and KC with WT mice (Supplemental Figure S5A and B; detailed gene list can be found in Supplemental Excel Table S4). To determine the contribution of Aqp5 to the development and growth of neoplastic ducts, especially DBA^-^/Agr2^+^ terminal ducts, we crossed KNGC mice with whole body Aqp5 knockout mice (49) to generate KNGCA mice (Figure 6A). Knockout of Aqp5 in 4-week old KNGCA mice was validated by qRT-PCR (Figure 6B). Although, Aqp5 staining was highly expressed in DBA^-^ ductal cells from KNGC mice, it was diminished in KNGCA mice (Figure 6C). Histological examination revealed that the knockout of Aqp5 in KNGC mice significantly decreased the development and expansion of neoplastic ducts, when compared to similarly aged KNGC mice (Figure 6D and E). Moreover, KNGCA mice showed restored expression of acinar markers and reduced expression of Agr2 and CFTR (Figure 6F). Consistent with the expression data, KNGCA mice developed a higher percentage of amylase^+^ acinar cells than KNGC mice (Supplemental Figure S5C and D). To further characterize the impact of Aqp5 deletion on the development of Agr2^+^ ducts and its effect on proliferation, serial sections of pancreas from KNGC and KNGCA mice were stained using a combination of Agr2 or pS10HH3 with DBA-FITC. As shown in Figure 6G, the mitotic marker, pS10HH3, was mainly localized in DBA^-^/Agr2^+^ ducts in KNGC mice (Figure 6H). However, KNGCA mice showed decreased pS10HH3+ cells in DBA^-^ ducts, as well as a reduction in the percentage of Agr2^+^ ducts (Figure 6H). Taken together, these data reveal an important role of Aqp5 in the development and growth of the Agr2^+^ terminal ductal population in KNGC mice.

**Figure 6.**
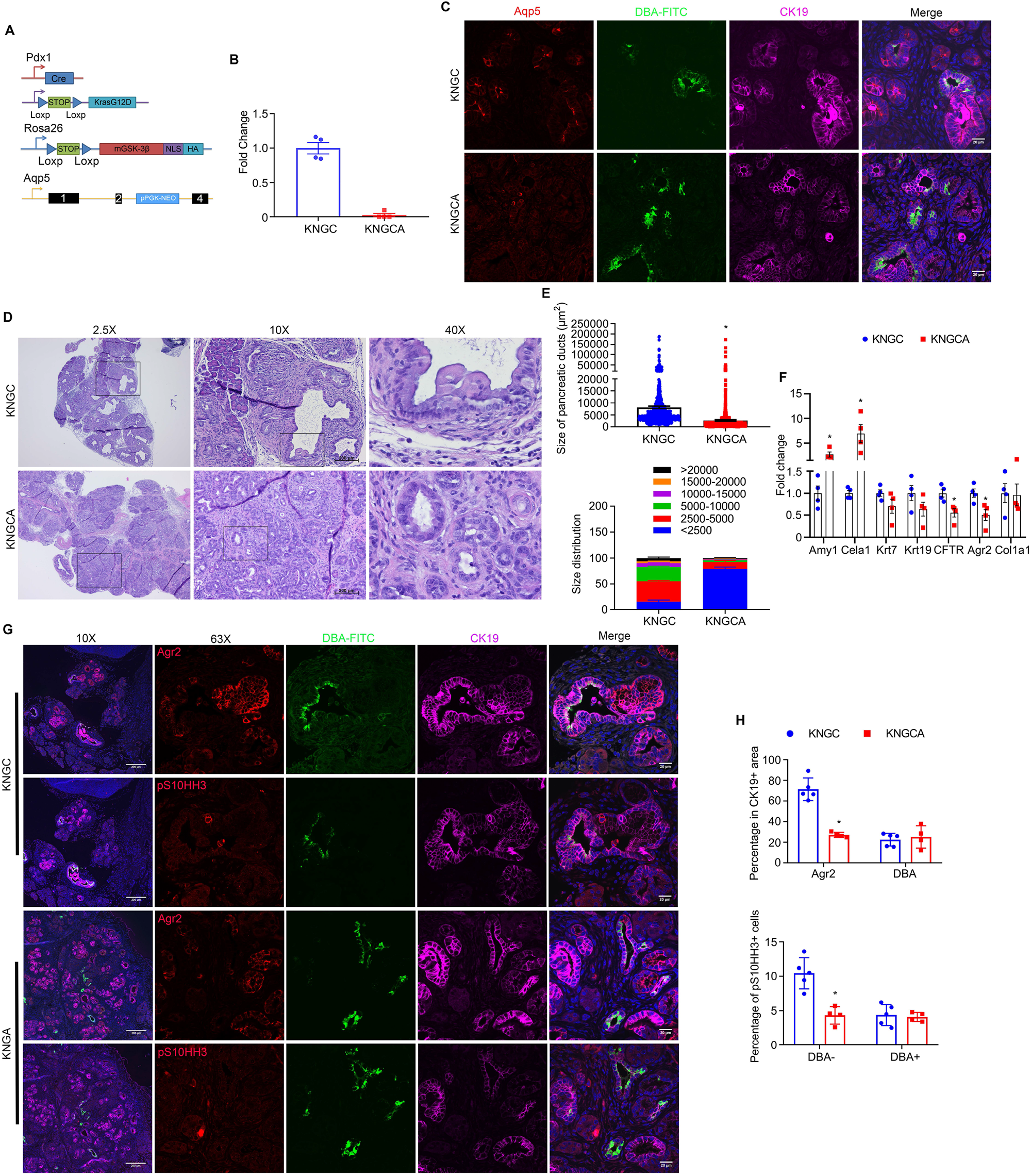
Aqp5 is necessary for the differentiation and growth of terminal ducts in KNGC mice. (A) Schematic representation of KNGCA (LSL-KRasG12D/Rosa26-LSL-nuclear GSK-3β/Pdx1-Cre/Aqp5 knockout) mouse model. Blue triangles indicate loxP sites. Black rectangles indicate exons of *aqp5* gene. (B) Real-time PCR quantification of Aqp5 gene expression from 4-week old KNGC and KNGCA mice. TBP, β-actin, RPLP0 and GAPDH were used as internal housekeeping gene controls. Data were analyzed and expressed as mean ± SEM. *P<0.05 KNGCA versus KNGC mice. (C) Immunofluorescence staining of Aqp5 (red), DBA-FITC (green) and CK19 (purple) from pancreatic sections of 4-week old KNGC and KNGCA mice. Nuclei were counterstained with Hoechst (blue). (D) H&E stained pancreatic sections from KNGC and KNGCA mice. Black boxes indicated area magnified. (E) Quantification of pancreatic ducts size distribution and average pancreatic duct size were analyzed and expressed as mean ± SEM. n = 3. *P<0.05 KNGCA versus KNGC mice. (F) Real-time PCR quantification of the indicated genes expressions from 4-week old KNGC and KNGCA mice. TBP, β-actin, RPLP0 and GAPDH were used as internal housekeeping gene controls. Data were analyzed and expressed as mean ± SEM. *P<0.05 KNGCA versus KNGC mice. (G) Immunofluorescence staining of Agr2 (red, upper panel) or pS10HH3 (red, lower panel) with DBA-FITC (green) and CK19 (purple) from serial pancreatic sections of 4-week old KNGC and KNGCA mice. Nuclei were counter-stained with Hoechst (blue). (H) Quantification of percentage in CK19 positive cells, as well as percentage of pS10HH3 positive cells in DBA positive and negative ducts were analyzed and expressed as mean ± SEM. *P<0.05 KNGC mice versus the other genotypes.

### Differences in Agr2, Aqp5 and DBA lectin staining in human IPMN

To investigate the distribution of Agr2^+^, Aqp5^+^ and DBA^+^ ducts in human IPMN, we performed Agr2/DBA-FITC IF staining and Aqp5 IHC staining on serial sections from tissue microarrays containing 140 human IPMN cases (see patient demographics of TMA in Supplemental Table S5) (50). Consistent with our KNGC mouse model, the majority of IPMN cores showed positive staining of both Agr2 and Aqp5 (Core 1 and 2 in Figure 7A and B). In contrast, we observed significantly fewer cores that were DBA^+/^Agr2^-^ (area 4 from Core 2 in Figure 7A). The strong and widespread distribution of Agr2^+^/Aqp5^+^/DBA^-^ terminal ducts in the IPMN TMA were further confirmed by Hscore, as Agr2 and Aqp5 staining were much higher than the DBA-FITC staining (Figure 7C). Moreover, correlation analysis of Agr2 and Aqp5 using Hscore indicated a statistically significant correlation between Agr2 and Aqp5 in the overall samples and in the subset of IPMN with adenocarcinoma (Figure 7D). To further test the significance of Agr2^+^/Aqp5^+^/DBA^-^ terminal ducts in IPMN development, we stained Agr2, Aqp5 and DBA in a previously published mouse model of IPMN, which combined mutations of KRas and GNAS (KGC) (51). Similar to the KNGC mice, IPMN lesions in the pancreas from KGC mice had higher staining for both Agr2 and Aqp5 than DBA-FITC (Supplemental Figure S6A and B). Collectively, these results support the conclusion that Agr2^+^/Aqp5^+^/DBA^-^ terminal ducts serve as a cell-of-origin for human IPMN lesions.

**Figure 7.**
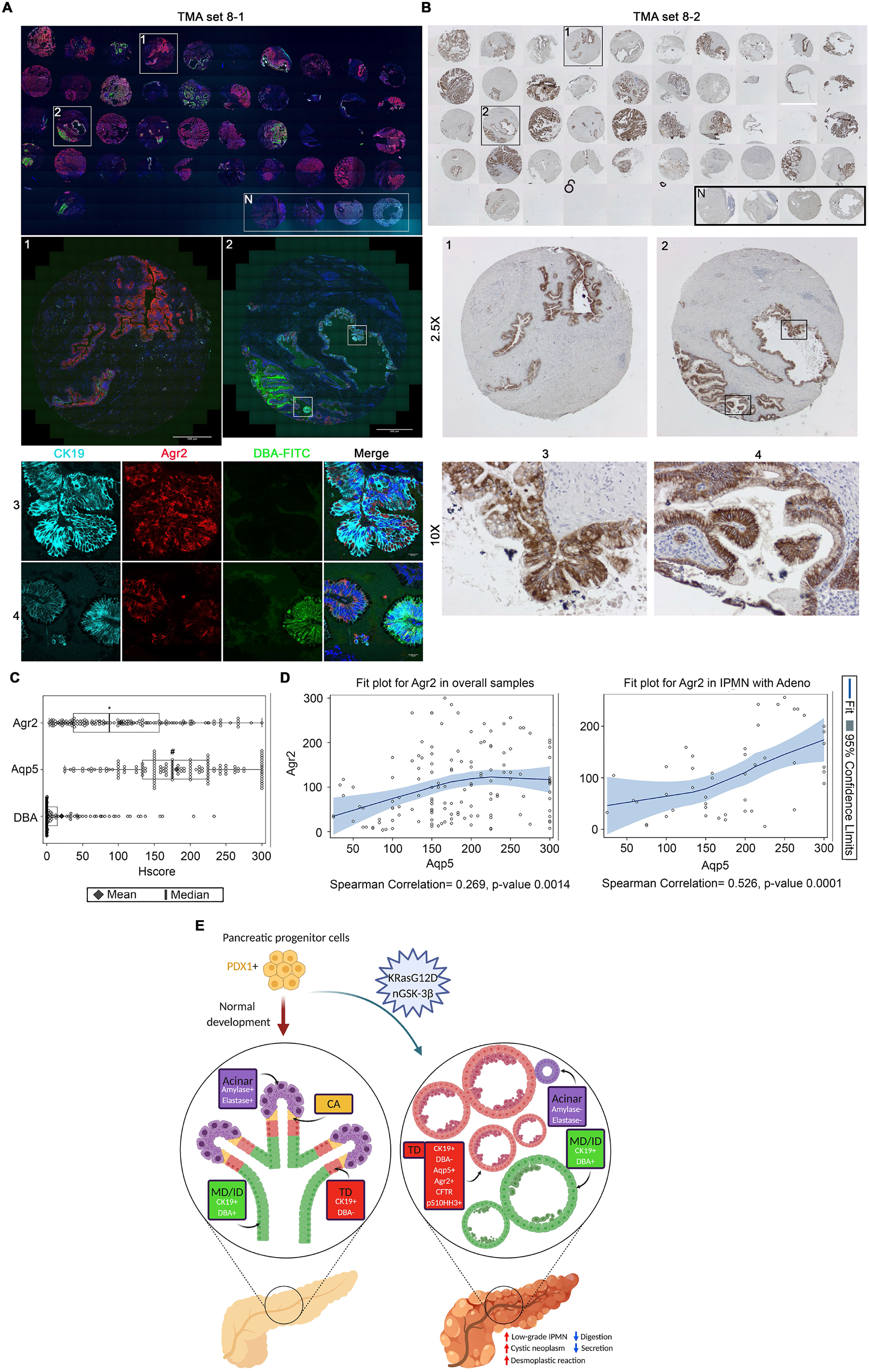
Agr2 and Aqp5, but not DBA staining is enriched in human IPMN. Immunofluorescence staining of Agr2 (red), DBA-FITC (green) and CK19 (Cyan) (A) and immunohistochemistry staining of Aqp5 (B) from serial sections of a TMA containing human IPMN. White boxes indicated area magnified. Nuclei were counterstained with Hoechst (blue) in (A). N: Normal control. (C) Histological score (Hscore) of Agr2, Aqp5 and DBA staining in 140 samples of human IPMN were calculated and graphed. Shown was mean ± SD. Black diamond: Mean value. Black line: Median value. *P<0.05 Agr2 versus DBA. #P<0.05 Aqp5 versus DBA. (D) Scatter plot with loess fit line and 95% confidence limits (colored area) for Agr2 in overall samples (Left) and IPMN with adenocarcinoma (Right) were drawn and Spearman correlation coefficient were calculated with P-value. (E) Schematic summary of terminal ducts as cell-of-origin of IPMN. PDX1^+^ pancreatic progenitor cells develop into normal pancreas (left) or low-grade IPMN with visible cysts in KNGC mice (right). Histological examination shows various markers described for each colored compartment of exocrine pancreas tissue including acinar (purple), centroacinar (CA, yellow), terminal ducts (TD, red) and main ducts/interlobular ducts (MD/ID, green).

## Discussion

In this study, we have generated a novel pancreas-specific nuclear GSK-3β expression and oncogenic KRas mouse model to investigate the contribution of nuclear GSK-3β to oncogenic KRas-driven pancreatic cancer development. Surprisingly, we found that nuclear GSK-3β rapidly promotes pancreatic cyst development and expansion of ductal cells in 4-week old KNGC mice. In a longitudinal study of KNGC mice, we observe notable reduction of body weight, as well as shortened overall survival, which is likely a result of the dramatic loss of pancreatic acinar cells. RNA-seq of the whole pancreas identified robust expression of Agr2, Aqp5 and CFTR, which are markers of PDA precursors (37,52). Consistent with previous publications (42,45,48), these markers are enriched in intercalated/terminal ducts and possess a higher proliferative potential than DBA-lectin^+^ interlobular ducts (Figure 7E). Significantly, knockout of Aqp5 in KNGC mice impaired IPMN development, indicating that Aqp5 is likely involved in terminal duct cell expansion/proliferation. We also observed the existence of widespread Agr2^+^/Aqp5^+^/DBA^-^ ducts in human IPMNs samples, and in a mouse model for IPMN (51). Taken together, these results suggest that terminal duct cells are a likely cell-of-origin for IPMN.

Several distinct precursor lesions, including PanIN, MCN and IPMN, are associated with the development of PDA. It is appreciated that KC mice develop PDA through a conversion of acinar-to-ductal metaplasia leading to PanIN lesion formation and PDA with long latency, which can be accelerated by the deletion or activation of mutant tumor suppressors, such as p16 or TP53 (9,10,14,15). Interestingly, using the same KC model, along with the addition of other oncogenes or tumor suppressors such as SMAD4, GNAS, Brg1, or Notch2, can result in a different progression path including the development of IPMN and MCN (51,53,54), reviewed in (15). It was first demonstrated by von Figura that Brg1 deletion using the *Ptf1a-Cre* promoter, which has Cre activity in almost all acinar cells and a subset of duct cells, promoted KRas-driven neoplasia in adult duct cells. However, using an inducible *Ptf1a*-CreER, in which Cre activity is restrained to mature acinar cells, they instead found Brg1 loss protected adult acinar cells from KRas-dependent PanIN development. Their results suggest that ductal cells are responsible for the IPMN-PDA progression in the *Ptf1a-Cre;* LSL-KRas; Brg1^f/f^ mouse model (54). Consistent with this study, Taki found that a *Ptf1a-Cre* inducible GEMM harboring GNAS^R201H^ and KRas^G12D^, two common somatic mutations found in human IPMN, cooperated to promote papillary dysplastic epithelia within dilated ducts mimicking human IPMN (53). In contrast, using inducible *Ptf1a-Cre;* GNAS^R201C^; KRas^G12D^ (KGC) showed a similar ability of acinar cells to give rise to IPMN, suggesting that it is actually underlying mutations, rather than the cell-of-origin, that are more important in IPMN development (51). Interestingly, using DBA to isolate pancreatic ductal cells from oncogenic KRas only or KRas; Brg1^f/f^ mice (37), showed that Brg1 sustained mature duct cell identity in the context of oncogenic KRas, whereas loss of Brg1 led to dedifferentiation of ductal cells with higher expression of progenitor markers (55). Thus, the cell of origin in these models still remains controversial. Although the lectin-binding protein DBA has been proposed to label the entire pancreatic ductal tree, including centroacinar cells, as well as PanIN and IPMN lesions, DBA low or negative duct structures were also observed in the previous studies (43,55). These DBA-negative structures were recognized as intercalated/terminal ducts, which possess stem cell or progenitor cell characteristics (9,43,48).

In the present study, we found that expression of nuclear GSK-3β and KRasG12D, under control of the Pdx1-cre transgene, which is required for differentiation of all pancreatic lineages, substantially initiates ductal expansion as early as the neonatal stage. Gene expressing profiling of different mucins from our RNA-Seq analysis showed elevated expression of muc1, muc5ac and muc6, which is consistent with the gastric type of IPMN (13). In addition, RNA-Seq data also revealed increased expression of Agr2, which is a recently identified marker used to distinguish ductal-derived PDA from acinar-derived PDA (41), thus suggesting that ductal cells are the main source for dilated ducts in KNGC mice. Moreover, expression of Agr2, together with Aqp5 and CFTR, three genes which have been demonstrated to have expression in human terminal ducts (40,45), were highly enriched in the DBA negative population from KNGC mice, further implicating terminal ducts as the cell-of-origin for IPMN development in the KNGC model. Significantly, KNGC mice showed a higher percentage of DBA^-^ cells undergoing mitosis as compared to DBA^+^ ducts. Lastly, the existence of Agr2^+^/Aqp5^+^ ductal cells as the prominent ductal cell population observed in human IPMN specimens, and the KGC mouse model, highlight terminal duct cells as the likely cell of origin in IPMN initiation.

Our previous studies, showing that deletion of GSK-3β in the KC model limits pancreatitis-induced acinar-to-ductal metaplasia, is in line with the data presented here, where genetic deletion of GSK-3β abrogates PanIN lesion development in aged KC mice. Taken together, these results suggest a cooperation between oncogenic KRas signaling pathways and GSK-3β signaling pathways leading to the development of PDA. In fact, we have previously shown that KRas signaling promotes GSK-3β gene expression, suggesting a possible feed-forward signaling loop (26). Indeed, we have found that GSK-3β becomes progressively over-expressed in human PDA and nuclear accumulated in the most aggressive tumors (24). While it is not clear what leads to the nuclear accumulation of GSK-3β, our data shown here would suggest that nuclear GSK-3β (comparing KNG to KNGC mice) alone cannot promote neoplastic transformation of ductal cells, but requires the cooperation of oncogenic KRas signaling. Although the mechanism by which these two signaling pathways are promoting the expansion of terminal duct cells as early as the neonatal stage is not known, it is well appreciated that both GSK-3β and MEK/ERK signaling are involved in stem cell biology, specifically the regulation of proliferation and maintenance of stem cell identity (56). Interestingly, activation of oncogenic KRas and nuclear GSK-3β using the inducible Rosa26CreERT or Krt19CreERT mice failed to reproduce the IPMN phenotype in KNGC mice, suggesting a differentiation reprogramming in terminal ducts encoded by nuclear GSK-3β and oncogenic KRas signaling occurs only during pancreas embryonic development. It is established that Krt19CreERT does not produce Cre recombinase activity in the intercalated/terminal ducts (57), thus future studies using an inducible mouse model to activate Cre in the terminal duct compartment, such as SOX9CreERT are warranted (48,58).

In summary, our study provides evidence that terminal ducts serve as the cell-of-origin for human IPMN. Future studies using this model combined with the addition of tumor suppressor mutations will be of significant interest as it pertains to IPMN-PDA development. In addition, further characterization of the KNGC mouse model has the potential to identify clinical biomarkers in terminal ducts, which could be used as novel early detection tools for human PDA.

## Materials and Methods

### Generation of nuclear GSK-3β conditional knock-in mice and mouse lines

Conditional GSK-3β knock-in mice containing an SV40 nuclear localization sequence (NLS; nGSK-3β) and HA-tag were generated by the Transgenic and Gene Knockout Mouse Core at the Mayo Clinic according to established protocols (59). The nGSK-3β targeting construct was generated using the previously described Rosa26-targeting vector pR26-CAG/EGFP-Asc, which contains a CAG promoter, and loxP flanked Neo/Stop cassette. LSL-KrasG12D and Pdx1-Cre mice have previously been described (27,28). Pancreas specific overexpression of nuclear GSK-3β with oncogenic KRas mutation mice was generated by crossing Pdx1-cre/LSL-KRasG12D (KC) mice with LSL-nuclear GSK-3β (NG) mice to produce Pdx1-cre/LSL-KRasG12D/LSL-nuclear GSK-3β (KNGC) animals. Krt19-Cre^ERT^ mice were obtained from the Jackson Laboratory (Stock Number: 026925). Rosa26-Cre^ERT^ mice were a gift from Dr. Hu Zeng (Mayo Clinic, MN, USA) and Aqp5 knockout mice were obtained from Dr. Varadaraj Kulandaiappan (Stony Brook University, NY, USA). These lines were intercrossed to obtain the desired genotypes used in this study. The data presented are from mice that have been backcrossed onto C57/B6 mice for 6 to 10 generations. Mice were housed in a barrier facility, and all experiments were performed with littermate-matched pairs, both male and female. All procedures were carried out according to the guidelines from and were approved by the Mayo Clinic Institutional Animal Care and Use Committee.

### Tamoxifen induction and acute pancreatitis model

Cre expression and activity was induced in 6-8 week old Krt19-Cre^ERT^ and Rosa26-Cre^ERT^ mice by intraperitoneal injection of tamoxifen (Sigma) dissolved in corn oil (Sigma) at a dose of 100 mg/kg of body weight. For mice carrying Krt19-Cre^ERT^, tamoxifen was injected once a day for 3 days, and for mice carrying Rosa26-Cre^ERT^, once a day for 5 days. To induce pancreatitis, 6-8 week old sibling littermates from Pdx1-cre (WT) and Pdx1-cre/nuclear GSK-3β (NGC) were selected for treatment. Mice were fasted for 12 h and allowed water ad libitum one day prior to the experiment. Animals were injected intraperitoneally into the right lower quadrant with a 50 μg/kg/bw (body weight) of caerulein dissolved in 0.9% saline in a volume of 100 μl. Injections were given at hourly intervals up to 8 times. Pancreata were harvested 7 days after caerulein treatment.

### Pancreatic ductal cell isolation

Preparation of single-cell suspensions from the mouse pancreas and pancreatic ductal cell isolation was performed as described (37,38,60). Briefly, mouse pancreata were removed and soaked overnight in cold 0.25% trypsin-EDTA (Thermofisher Scientific). The upper enzymatic fluid was aspirated and the digestion was continued for 10 min at 37°C with residual trypsin-EDTA and DNase I (Sigma, 10 μg/ml) in 3 ml of DMEM/F12 medium (Thermofisher Scientific). The pancreas tissue was further digested in DMEM/F12 supplemented with collagenase D (Sigma, 2.5 mg/ml) and liberase DL (Sigma, 0.5 mg/ml), and DNase I for 10 min at 37°C with gentle shaking. The digestion was terminated and quenched by excess DMEM/F12 supplemented with 1X GlutaMAX (Gibco), 10 mmol/L HEPES (Gibco), and 1% of penicillin/streptomycin (Gibco). Cells were gently pipetted to maximize the release of single cells and spun down, and resuspended in ACK lysis buffer to eliminate red blood cells. ACK buffer was quenched with 2% fetal bovine serum (FBS) in PBS and cells were resuspended in a sorting buffer containing PBS, 0.5% bovine serum albumin, and 2 mM EDTA, followed by 10 min incubation in fluorescein-labeled *Dolichos biflorus agglutinin* (DBA) lectin (Vector Laboratories) with agitation at 4°C. Cells were washed in the sorting buffer and resuspended in the same buffer with anti-FITC Microbeads (Miltenyi Biotec), and incubated on a rotor for 15 min at 4°C. Separation was performed using MS columns (Miltenyi Biotec), according to the manufacturer’s protocol.

### Immunoblot Analysis

Collected pancreata or isolated pancreatic ductal cells were lysed with Western lysis buffer (1% Triton X-100, 10 mM Tris Base, 50 mM NaCl, 5 mM EDTA, 50 mM NaF, 30 mM Na4P2O7 pH 7.4) supplemented with aprotinin, leupeptin, sodium orthovanadate, phenylmethylsulfonyl fluoride (PMSF) and calyculin A (Cell Signaling Technologies, Beverly, MA, USA). Lysates were subjected to sodium dodecyl sulfate (SDS)-polyacrylamide gel electrophoresis and immunoblotting as described (27). Antibodies used for immunoblotting and immunofluorescence are described in detail in Supplemental Excel Table S1.

### IHC, EdU labeling and Immunofluorescence (IF)

Mice were anesthetized using isoflurane (Nova Plus Pharmaceuticals), followed by cervical dislocation. The whole pancreas was quickly removed and fixed overnight in 4% PFA with gentle shaking, embedded in paraffin, cut into 5 μm-thick sections. Sections were subjected to H&E, IHC and immunofluorescence staining as described (27). Briefly, sections were de-paraffinized and hydrated by graded washes with xylene and ethanol. Antigen retrieval was accomplished by sub-boiling in 10 mM sodium citrate acid buffer (pH 6.0) for 15 min. Endogenous peroxidase activity was blocked by incubation of slides in 3% H_2_O_2_ for 10 min. Nonspecific binding was blocked with 5% normal goat serum. Sections were then incubated with primary antibodies overnight at 4°C. Anti-rabbit secondary (#8114, Cell Signaling Technologies, Beverly, MA, USA) and diaminobenzidine substrate kit (#8059, Cell Signaling Technologies, Beverly, MA, USA) were used for immunohistochemistry. Slides were then counterstained with Mayer’s Hematoxylin before dehydration and mounting. For immunofluorescence, slides were incubated with fluorescent-conjugated secondary antibodies for 1 h at room temperature before confocal scanning. For EdU labeling, mice were injected with EdU at a concentration of 50 mg/kg/bw in saline 2 hours before sacrifice. Staining was performed by Click-iT^®^ EdU Alexa Fluor^®^ 647 Imaging Kit following manufacturer’s instructions (Thermofisher scientific, USA). Confocal images were collected with an LSM-800 laser scanning confocal microscope with a 63x-oil Plan-Apochromat objective lens using ZEN Blue 2.6 software package (Carl Zeiss, Oberkochen, Germany). For whole pancreas tissue section scanning, stage marks were placed around the edge of the pancreas and tile region was drawn to cover all the stage marks. Images were taken under 10x objective lens using tiles. The stitching method within the Zen Blue software package was applied to process the tile images into one final image. The percentage of EdU-647 + cells was enumerated, and the area and integrated density of indicated staining were measured using the ImageJ open source image-processing package.

### RNA isolation and quantitative RT-PCR

RNA isolation and quantitative RT-PCR were performed as previously described (27). Briefly, pancreatic total RNA or isolated pancreatic ductal cells was isolated using Trizol and further purified with an RNeasy Mini Kit (Qiagen, Valencia, CA). Reverse transcription was performed with the Superscript III RT-PCR Kit (Invitrogen). Quantitative PCR was performed with the SYBR Green PCR Master Mix using the ABI StepOnePlus Sequence Detection System (Applied Biosystems, Carlsbad, CA). Four housekeeping genes were used for normalization of gene expression. The double Δ Ct method was used to analyze gene expression. Experiments were performed a minimum of three times using independent cDNAs. Primer sequences are provided in Supplemental Excel Table S2.

### RNA-Seq and data analysis

Total RNA was isolated from 4 weeks old littermates as described above and the average RNA integrity number values were measured by an Agilent Bioanalyzer. Individual transcriptome sequencing (RNA-Seq) libraries were prepared from each mouse using an Illumina TruSeq v2 kit. 100 base pairs reads were collected with an Illumina HiSeq 4000 instrument. Fastq files of paired-end reads were aligned with STAR 2.6.0a (61) to the UCSC reference genome mm10 with basic 2-pass mapping. Gene counts were obtained using the subRead featureCounts program 1.4.6 (62) based on the UCSC mm10 annotation. Differential expression analyses were performed using R package DESeq2 1.10.1 (63) after removing genes with average raw counts less than 10. Genes with log2 fold change greater than 2 or less than −2, and FDR less than 0.05 were considered significantly differentially expressed. For functional annotation analysis, the Fisher’s exact test was used to determine overrepresented pathways or gene sets in significantly up- or down-regulated genes, against gene sets described in Enrichment Map (64) and R package KEGG.db (65). RNA sequencing data have been deposited in the Gene Expression Omnibus under the accession number GSE153548.

### Staining on Tissue Microarrays

The tissue microarrays (TMAs) consisted of 140 unique individuals with IPMN who were acquired from the Mayo Clinic SPORE in Pancreatic Cancer. IHC for Aqp5 and immunofluorescence staining of Agr2/DBA/CK19 were performed as above. Cores were excluded if absent in the slide. Aqp5 histological scoring was performed and evaluated by pathologist (Lizhi Zhang) as follows, staining intensities were scored from 0 (no staining) to 3 (high staining) and extent was scored from 0 (negative), 1 (<25%), 2 (25-50%), 3 (50-75%), to 4 (>75%, widespread staining). Whole slides scanning and measurement of intensity and area of Agr2/DBA/CK19 immunofluorescence staining in TMAs were described above and staining intensities were scored from weak (<33%), medium (33-66%) to strong (>66%) and extent was scored as the percentage of Agr2/DBA in CK19 positive cells. The H-score system was used for IHC evaluation of Aqp5 and IF evaluation of Agr2 and DBA by multiplying the extend or percentage of cells with staining intensity ordinal value ranging from 0 to 300. H-scores were available on 133 to 135 of the 140 patients represented on the TMA.

### Statistical analysis

Data are expressed as mean ± SEM and analyzed by repeated measures analysis of variance, one-way ANOVA and unpaired Student’s t-test using GraphPad Prism software (GraphPad Software Inc., La Jolla, CA). A value of p <0.05 denotes statistical significance. Scatterplots including a loess smoother (smoothing parameter=0.8) were generated to visualize the relationship between AGR2 and AQP5. Spearman correlation coefficient and p-value are reported to summarize the nature of the relationship.

## Supporting information

Excel File containing Supplemental Tables S1-S5

## Acknowledgement

We would like to thank members of the Division of Oncology Research especially Drs. Scott Kaufmann, Zhenkun Lou, Martin Fernandez-Zapico as well as members of the Billadeau laboratory for helpful discussions. We would like to thank Dr. Varadaraj Kulandaiappan (Stony Brook University, NY, USA) for the Aqp5 knockout mice. Lastly, we would like to thank Dr. Howard Crawford and other members of the Cold Spring Harbor Pancreatic Cancer Workshop – for thought-provoking discussions that helped shape this project.

