## Supplemental Figures 1-6 and Legends for "Nuclear GSK-3β and Oncogenic KRas Promote Expansion of Terminal Duct Cells and the Development of Intraductal Papillary Mucinous Neoplasm"

### Supplemental Figure S1

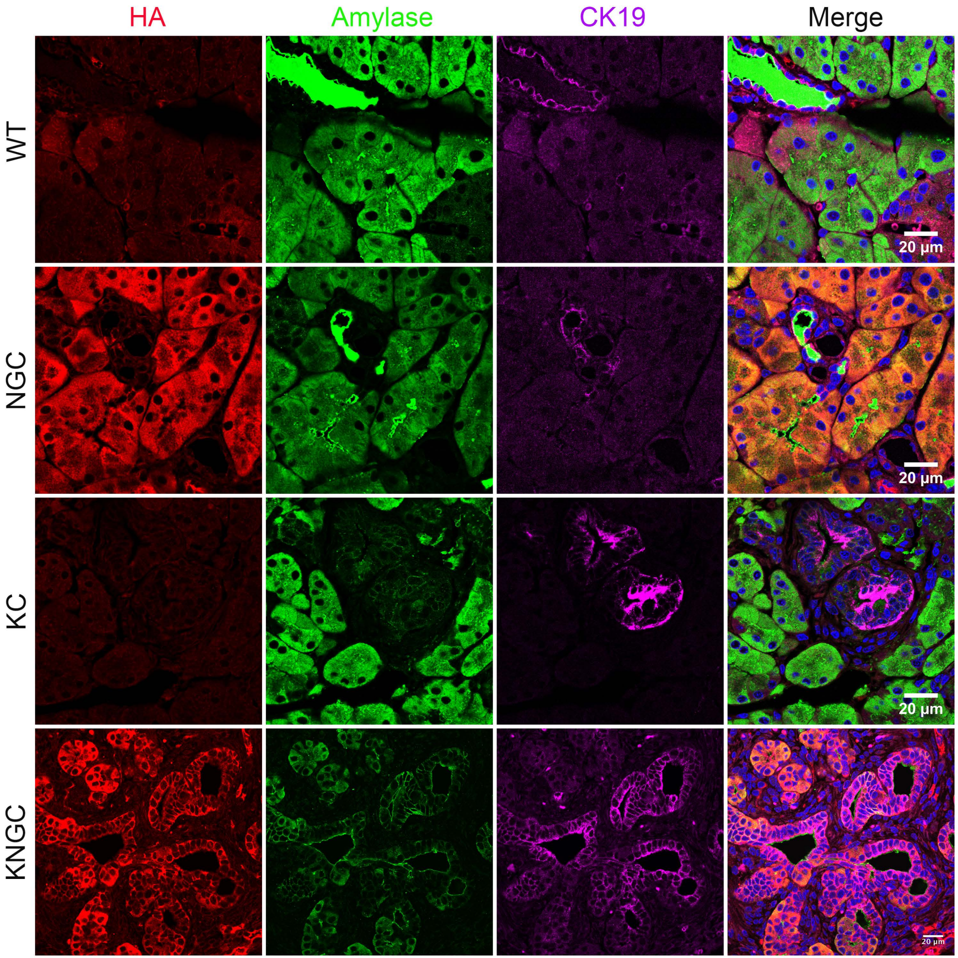

### Supplemental Figure S2

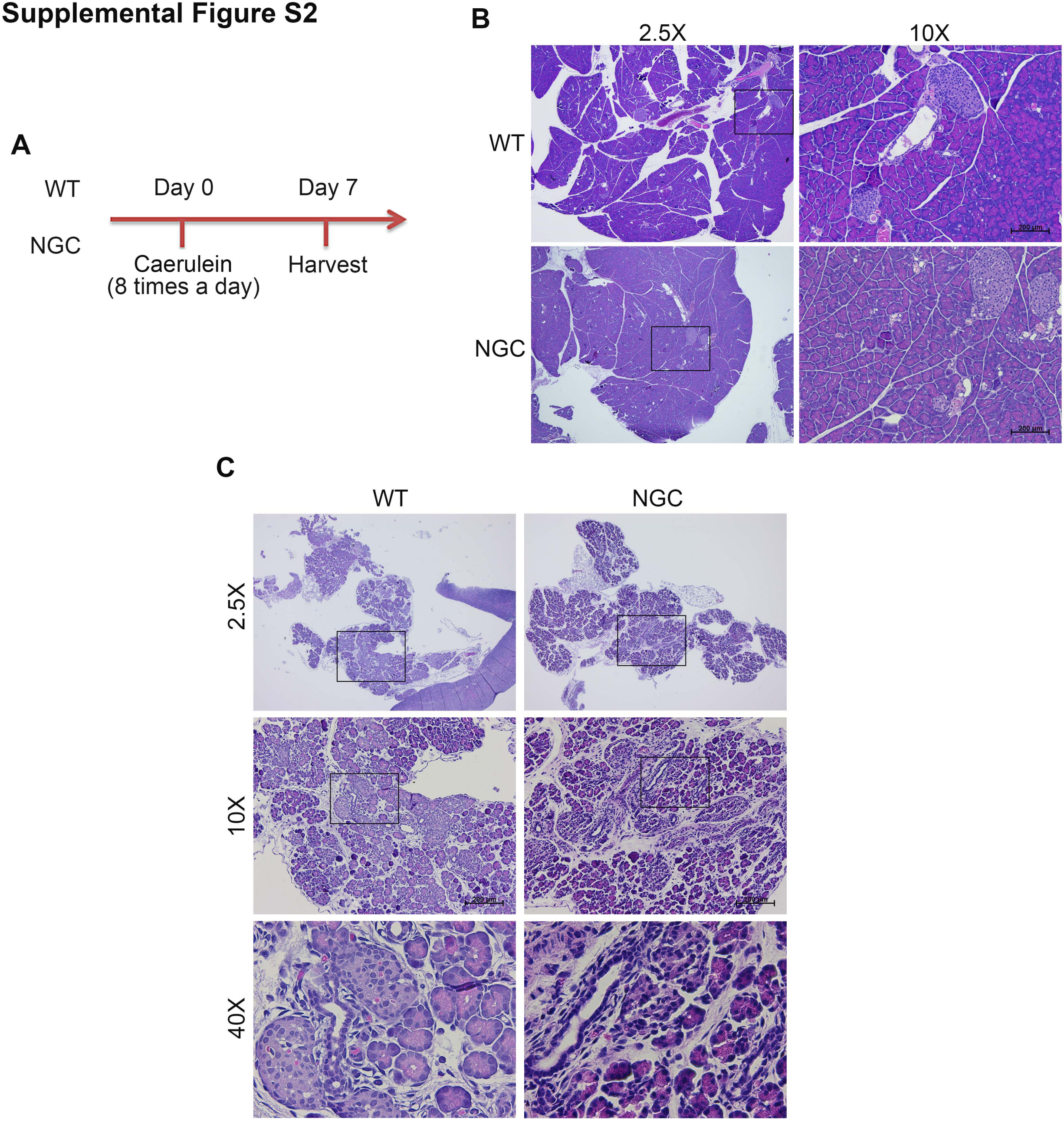

#### Supplemental Figure S3

**A**

| | NGC vs WT | KC vs WT | KNGC vs WT | KNGC vs NGC | KNGC vs KC |
|---|---|---|---|---|---|
| mRNAs increased | 118 | 127 | 5304 | 4474 | 5056 |
| mRNAs decreased | 48 | 73 | 4593 | 4603 | 4297 |
| Total | 166 | 200 | 9897 | 9077 | 9353 |

**B**

KEGG pathways with FDR<0.05

| Cutoffs | KNGC vs NGC | KNGC vs KC |
|---|---|---|
| $Log_2FC > 2$ | 1131 | 1491 |
| $Log_2FC > 3$ | 407 | 327 |
| $Log_2FC < -2$ | 122 | 191 |
| $Log_2FC < -3$ | 68 | 107 |

**C**

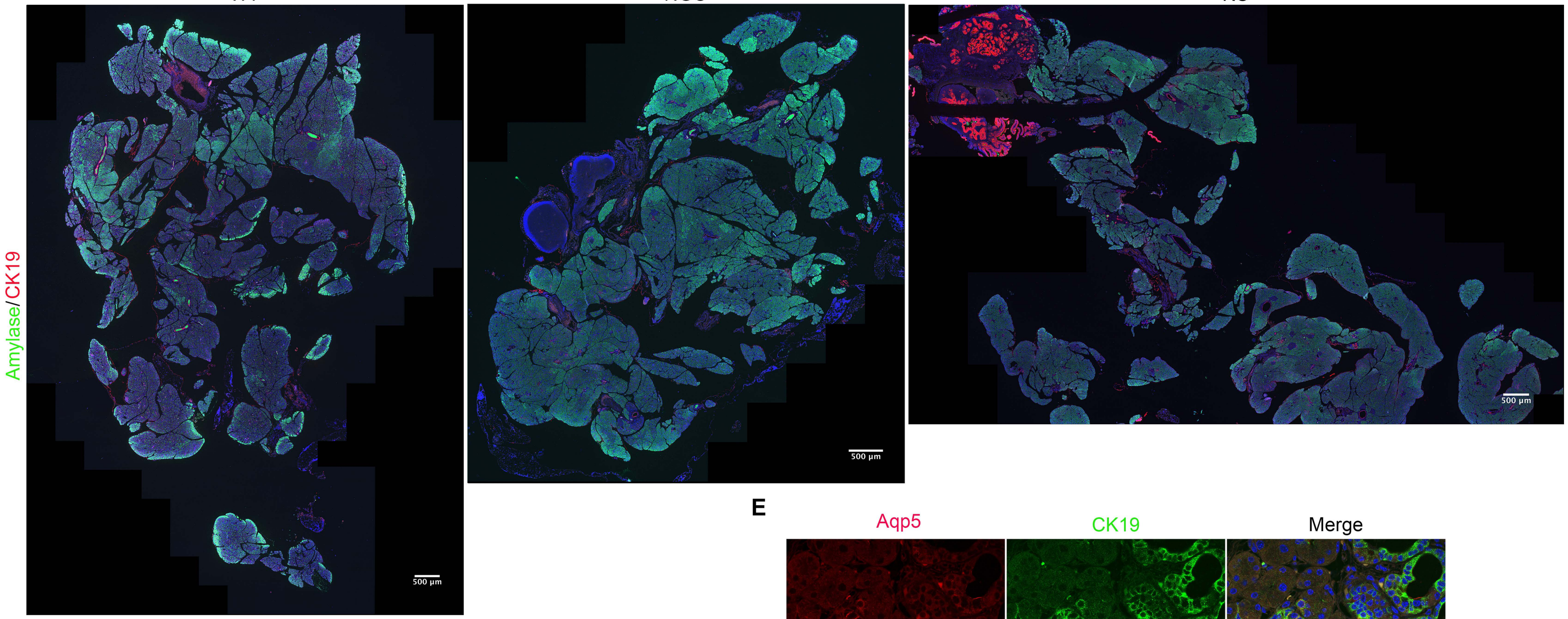


**D**

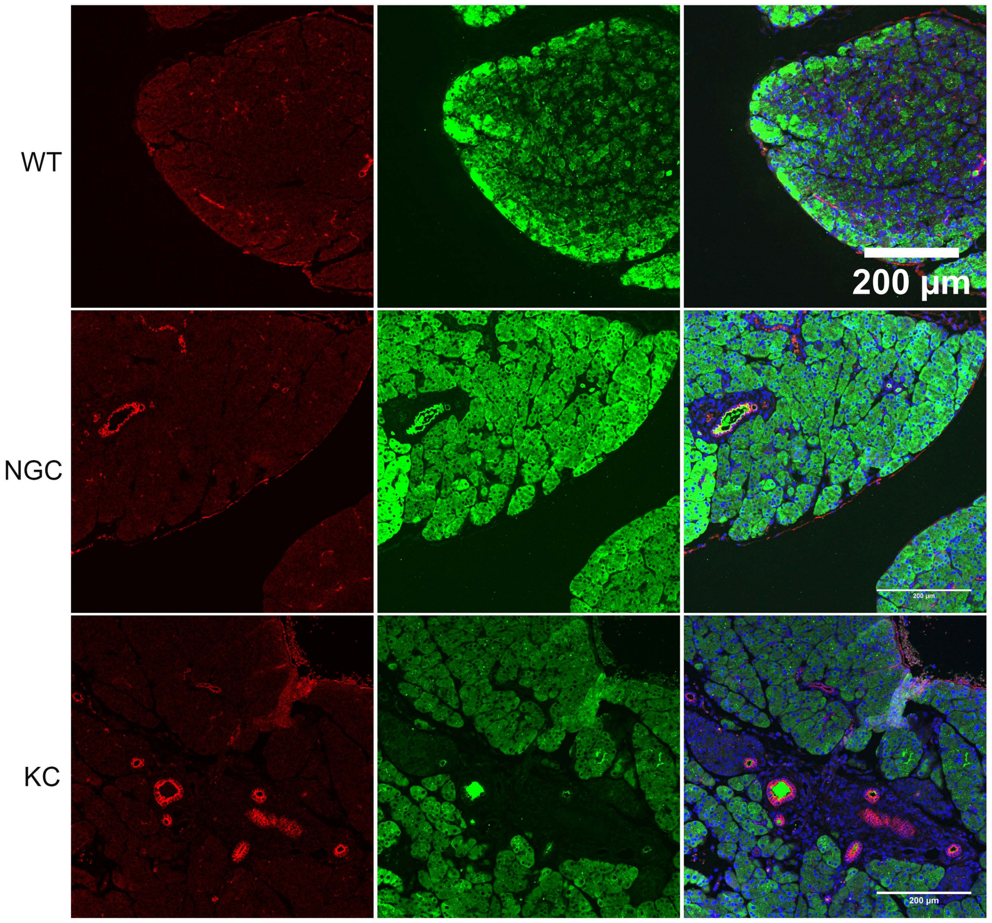


**E**

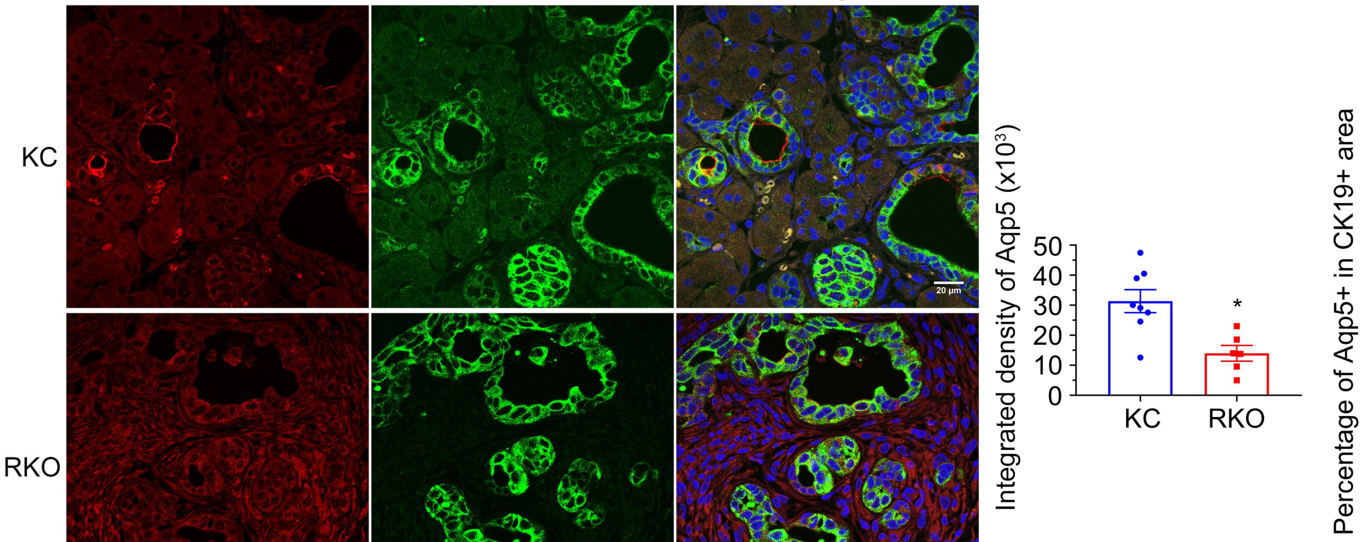


**F**

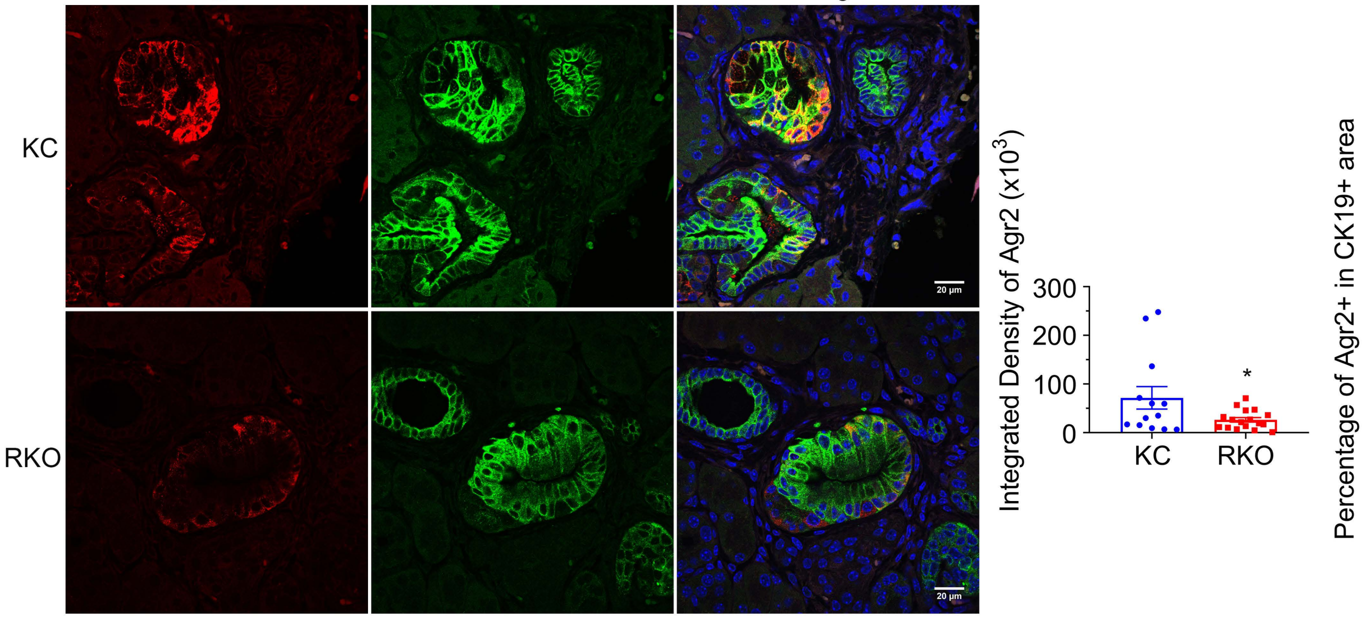

### Supplemental Figure S4

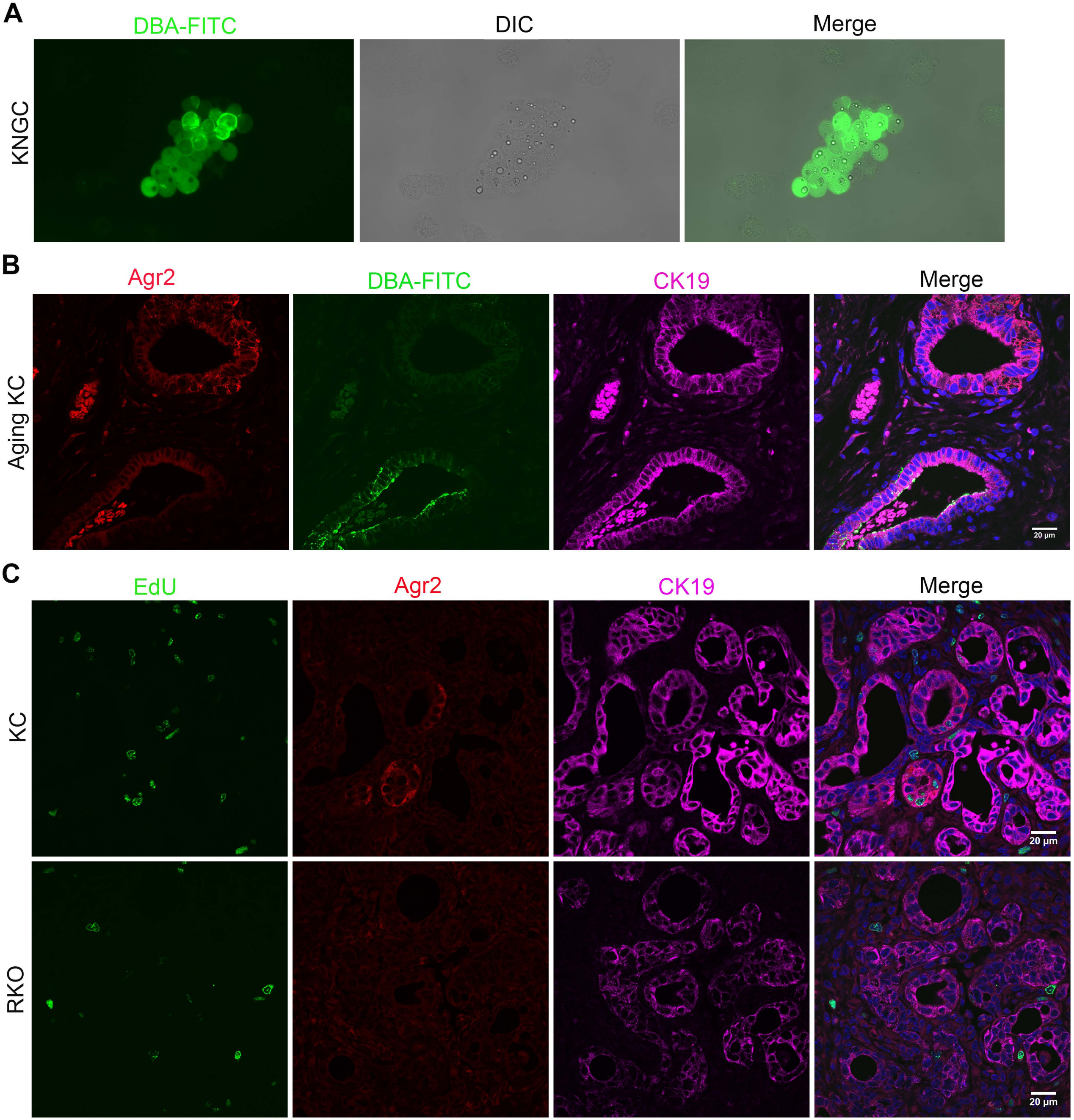

#### Supplemental Figure S5

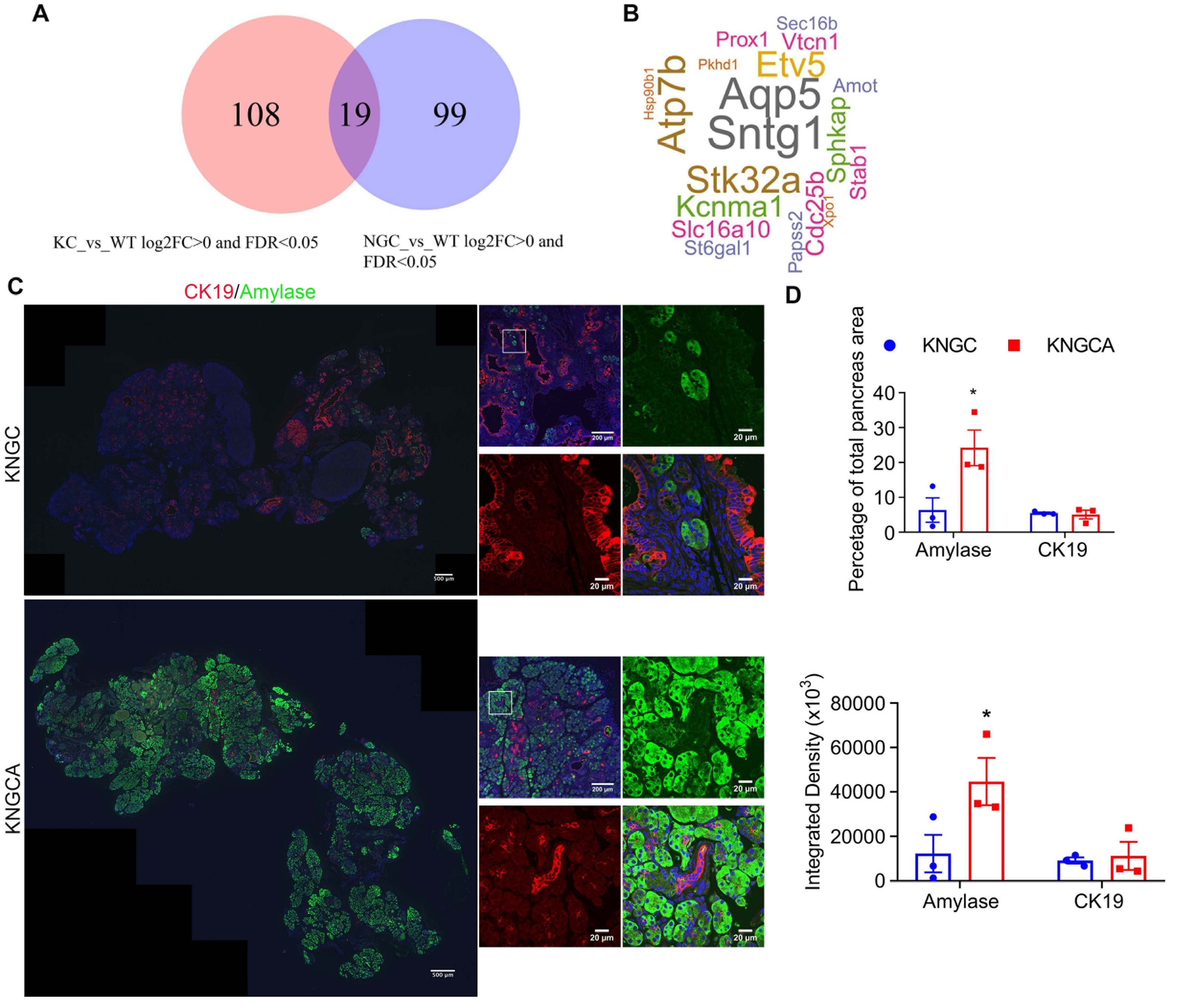

#### Supplemental Figure S6

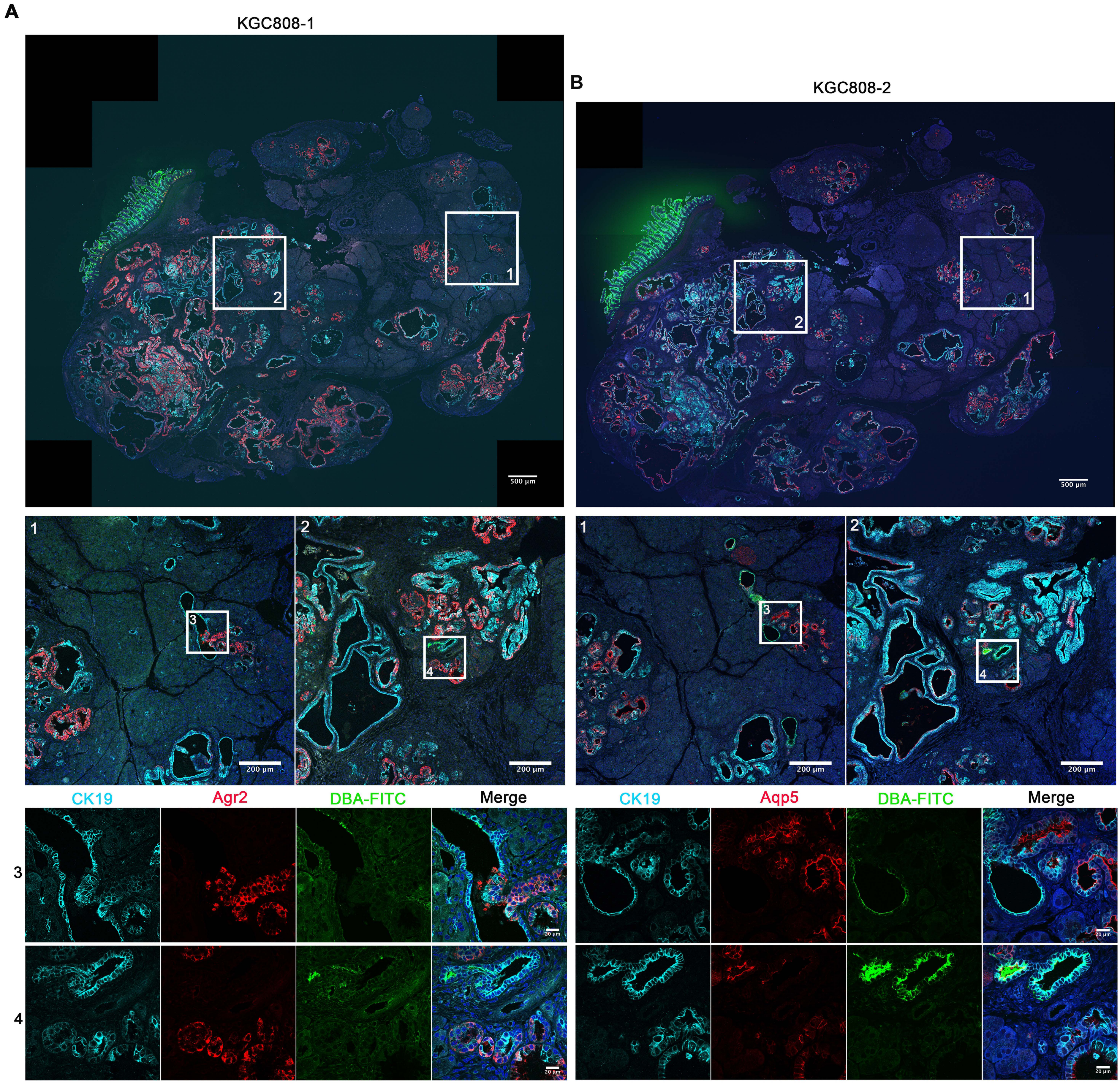

**Supplemental Excel File and Figures legend**

**Supplemental Excel File: Table S1 –** Antibodies used in this study, **Table S2 –** primers used for qPCR, **Table S3 –** complete list of group markers (Figure 4), **Table S4 –** commonly increased genes (Figure 6), and **Table S5 –** patient demographics for TMA (Figure 7).

**Figure S1. Nuclear GSK-3β and oncogenic KRas promote pancreatic ductal cells expansion and IPMN development.** Immunofluorescence staining of HA (red), amylase (green) and CK19 (purple) from pancreatic sections of indicated genotypes at 4 weeks age. Nuclei were counter-stained with Hoechst (blue). Shown are representative image from 3 different mice.

**Figure S2. Nuclear GSK-3β and oncogenic KRas initiate ductal hyperplasia at an early stage of pancreas development.** (A) Scheme for caerulein-induced acute pancreatitis model and analysis. (B) H&E stained pancreatic sections from WT and NGC mice treated with caerulein as indicated in (A). (C) H&E stained pancreatic sections from neonatal WT and NGC mice. Black boxes indicated area magnified.

**Figure S3. Transcriptional regulation of pancreatic ductal neoplasia by overexpression of nuclear GSK-3β and oncogenic KrasG12D activation.** (A) Differential expression analyses by DESeq2 of RNA-Seq quantification of mRNA changes from different comparison as indicated, with an FDR cutoff of <0.05. (B) Number of enriched KEGG signaling pathways with FDR<0.05 in comparison between KNGC and NGC or KC mice, different log2 fold change ($Log_2FC$) were used as cutoffs. (C) Whole slide scanning of immunofluorescence staining of Amylase (green) and CK19 (red) from pancreatic sections of indicated genotypes at 4-weeks of age. (D) Representative image of immunofluorescence staining of Amylase (green) and CK19 (red) from pancreatic sections of indicated genotypes at 4-weeks of age. (E) Immunofluorescence staining of Aqp5 (red) and CK19 (green) from pancreatic sections of aging KC and RKO mice as indicated in Figure 1. Quantification of Aqp5 positive percentage in CK19 cells and integrated density were analyzed and expressed as mean ± SEM. *P<0.05 RKO mice versus KC mice. (F) Immunofluorescence staining of Agr2 (red) and CK19 (green) from pancreatic sections of aging KC and RKO mice as indicated in Figure 1. Quantification of Agr2 positive percentage in CK19 cells and integrated density were analyzed and expressed as mean ± SEM. *P<0.05 RKO mice versus KC mice. Nuclei were counter-stained with Hoechst (blue).

**Figure S4. Expression of nuclear GSK-3β with oncogenic KRas leads to the development of two distinct ductal populations.** (A) Green fluorescent channel and differential interference contrast (DIC) image of isolated pancreatic cells from KNGC mice after DBA-FITC labeling and anti-FITC-conjugated microbead separation. (B) Immunofluorescence staining of Agr2 (red), DBA-FITC (green) and CK19 (purple) from pancreatic sections of aging KC mice as indicated in Figure 1. (C) Immunofluorescence staining of Agr2 (red), EdU (green) and CK19 (green) from pancreatic sections of aging KC and RKO mice as indicated in Figure 1. Nuclei were counter-stained with Hoechst (blue).

**Figure S5. Aqp5 is necessary for the differentiation and growth of terminal ducts in KNGC mice.** (A) Venn diagram showing the number of genes fitting the indicated cutoffs. (B) Word cloud using 19 common genes shown in (A). (C) Immunofluorescence staining of amylase (green) and CK19 (red) from pancreatic sections of aging KNGC and KNGCA mice at 4 weeks of age. White boxes indicate magnified area. Nuclei were counter-stained with Hoechst (blue). Shown are representative images. (D) Quantification of percentage and integrated density of Amylase and CK19 staining were analyzed and expressed as mean ± SEM. *P<0.05 KNGCA mice versus KNGC mice.

**Figure S6. Differences in Agr2, Aqp5 and DBA staining in human IPMN.** Immunofluorescence staining of (A) Agr2 (red) and (B) Aqp5 (red) with DBA-FITC (green) and CK19 (cyan) from pancreatic serial sections of KGC mice at 11 weeks post Dox induction. White boxes indicate magnified area. Nuclei were counter-stained with Hoechst (blue).
